# Aberrant landscapes of maternal meiotic crossovers contribute to aneuploidies in human embryos

**DOI:** 10.1101/2023.06.07.543910

**Authors:** Daniel Ariad, Svetlana Madjunkova, Mitko Madjunkov, Siwei Chen, Rina Abramov, Clifford Librach, Rajiv C. McCoy

## Abstract

Meiotic recombination is crucial for human genetic diversity and chromosome segregation accuracy. Understanding its variation across individuals and the processes by which it goes awry are long-standing goals in human genetics. Current approaches for inferring recombination landscapes either rely on population genetic patterns of linkage disequilibrium (LD)—capturing a time-averaged view—or direct detection of crossovers in gametes or multi-generation pedigrees, which limits dataset scale and availability. Here, we introduce an approach for inferring sex-specific recombination landscapes using data from preimplantation genetic testing for aneuploidy (PGT-A). This method relies on low-coverage (<0.05*×*) whole-genome sequencing of *in vitro* fertilized (IVF) embryo biopsies. To overcome the data sparsity, our method exploits its inherent relatedness structure, knowledge of haplotypes from external population reference panels, as well as the frequent occurrence of monosomies in embryos, whereby the remaining chromosome is phased by default. Extensive simulations demonstrate our method’s high accuracy, even at coverages as low as 0.02*×*. Applying this method to PGT-A data from 18,967 embryos, we mapped 70,660 recombination events with *∼*150 kbp resolution, replicating established sex-specific recombination patterns. We observed a reduced total length of the female genetic map in trisomies compared to disomies, as well as chromosome-specific alterations in crossover distributions. Based on haplotype configurations in pericentromeric regions, our data indicate chromosome-specific propensities for different mechanisms of meiotic error. Our results provide a comprehensive view of the role of aberrant meiotic recombination in the origins of human aneuploidies and offer a versatile tool for mapping crossovers in low-coverage sequencing data from multiple siblings.

## Introduction

Recombination between homologous chromosomes is a key source of human genetic diversity [1, 2]. The crossovers that mediate such genetic exchanges during meiosis are also important for ensuring the accuracy of chromosome segregation [3, 4]. Notably, female meiosis initiates during fetal development, when homologs pair, acquire double-strand breaks, and establish crossovers that form physical linkages (chiasmata) to stabilize the chromosomes. Such chiasmata must then be maintained over decades-long meiotic arrest, until meiosis resumes at ovulation. Abnormal number and/or location of crossovers may predispose oocytes to gains or losses of whole chromosomes (aneuploidies), which are the leading cause of human pregnancy loss and congenital disorders [5]. Hypotheses about the role of recombination in aneuploidy formation largely originated from studies of model organisms [4, 6, 7]. Meanwhile, the smaller number of studies in humans have primarily focused on the subset of trisomies that are compatible with *in utero* development [8, 9, 10, 11], with less focus on the most common trisomies (Chr 15, Chr 16, and Chr 22) observed in oocytes and preimplantation embryos (though see [12, 13, 14]). To overcome this limitation, several previous studies have analyzed all products of meiosis (i.e., the first and second polar body, as well as a biopsy of the corresponding embryo) [15, 16]. While insightful, such sampling is technically demanding, limiting sample sizes and in turn limiting power and resolution for comparing genetic maps.

Over the last decade, several studies on human embryos have been conducted within the framework of preimplantation genetic testing for monogenic disorders (PGT-M) using methods such as Karyomapping [17], siCHILD/haplarithmisis [18], OnePGT [19], and GENType [20], whereby parental DNA is assayed along with that of the embryos, and unaffected embryos are prioritized for transfer. These genome-wide haplotyping methods allow mapping of maternal and paternal crossovers along chromosomes. However, the number of patients that undergo PGT-M is small compared to the number of patients that undergo preimplantation genetic testing for aneuploidy (PGT-A), again limiting answers to broader questions about the crossover landscape. For example, a recent PGT-M study by Tšuiko et al. [21] inferred the parental and mechanistic origin of chromosome abnormalities in 2706 embryos from PGT-M patients and found 269 trisomies in total. A similar PGT-M study of recombination by Ma et al. [22] analyzed 1519 embryos and 353 autosomal aneuploidies.

Other current approaches for inferring the landscape of recombination rely either on population patterns of linkage disequilibrium (LD)—capturing a time-averaged view of historical recombination events—or direct detection of crossovers based on genotyping of haploid gametes or multi-generation pedigrees (e.g., parent-offspring trios), again limiting the scale and availability of relevant datasets [23, 24, 25, 26]. Moreover, most of these methods are designed for discovering recombination using data from normal, disomic chromosomes. Mapping meiotic crossovers in large samples of both normal and aneuploid embryos using a unified statistical framework would allow a robust test of the role of recombination in the genesis of aneuploidy.

To this end, we introduce a statistical approach tailored to sequencing data from PGT-A, which is based on low-coverage (*<*0.05*×* per homolog) whole-genome sequencing of biopsies from *in vitro* fertilized (IVF) embryos. We retrospectively apply our method to normal disomic chromosomes identified in existing low-coverage PGT-A data from 18,967 embryos and replicate features of sex-specific recombination maps that were previously described based on large prospective studies of living populations. We then extend our method to trisomies, testing the extent to which the landscape of recombination differs between normal and aneuploid chromosomes. Together, our study sheds light on the dual function of meiotic recombination in generating genetic diversity while ensuring fidelity of human meiosis.

## Results

### A method for inferring crossovers based on low-coverage sequencing data from multiple siblings

One general approach for discovering the genomic locations of meiotic recombination events is to compare genotype data from related individuals. Such data can be scanned to identify regions where haplotypes match (i.e., are identical by descent [IBD]). The boundaries of the matched haplotypes reflect the locations of meiotic crossovers in the history of the sample. The information gained by comparing haplotypes among relatives serves as the foundation for several different approaches for PGT-M [17, 18, 19]. However, directly calling diploid genotypes from sequencing data requires a minimum coverage of 2*×* (to sample both alleles) and in practice requires coverage several fold higher to overcome technical challenges such as coverage variability, ambiguous alignments due to repetitive sequences, and other sequencing and analytic artifacts. Because data from PGT-A typically fall well below these coverage requirements, they are generally assumed unsuitable for applications that demand genotypes, including the study of recombination landscapes in embryos. However, as exemplified by common methods such as genotype imputation [27], knowledge of patterns of LD from external population genetic reference panels may facilitate the extraction of meaningful signal from sparse, low-coverage datasets, including in the context of prenatal genetics [28, 29].

Building on this logic, we introduce a haplotype matching approach, named Linkage Disequilibrium-Informed <u>C</u>omparison of <u>H</u>aplotypes <u>A</u>mong <u>S</u>ibling <u>E</u>mbryos (LD-CHASE), tailored to DNA sequencing data from PGT-A. Most current implementations of PGT-A involve low-coverage high-throughput sequencing of trophectoderm biopsies from IVF embryos at day 5 or 6 post-fertilization, with the goal of prioritizing chromosomally normal (i.e., euploid) embryos for transfer to improve IVF outcomes [30]. PGT-A offers a unique source of genomic data from large numbers of sibling samples, as each IVF cycle typically produces multiple embryos, and often multiple IVF cycles are necessary in infertility treatment.

Disomic chromosomes of any two sibling embryos will possess discrete genomic intervals with different counts of matching haplotypes, and transition points between these intervals reflect the locations of meiotic crossovers. The occurrence of monosomy (or uniparental isodisomy [isoUPD], isolated to individual chromosomes or genome-wide [GW-isoUPD]) among a set of sibling embryos greatly simplifies this comparison, as the remaining chromosome is phased by default, facilitating discovery of sex-specific crossovers (i.e., originating during gamete formation in one of the two parents). Here, we leverage the common occurrence of chromosome loss to reveal the sex-specific landscapes of meiotic crossovers among a large sample of IVF embryos.

Briefly, LD-CHASE uses sparse genotypes obtained from low-coverage sequencing data to identify the locations of meiotic crossovers (Figures 1 and S1). At such coverages, direct comparison of haplotypes is not possible, as a small minority of the genome is covered by any sequencing reads, and positions of aligned reads from samples under comparison rarely overlap. We circumvent this challenge based on patterns of LD, whereby observations of a set of alleles from one sequencing read may provide indirect information about the probabilities of alleles at nearby, unobserved variant sites. This in turn informs the relative probability that a given pair of reads originated from identical homologous chromosomes versus from distinct homologous chromosomes, which we formalize using a likelihood framework (see Methods). Transitions between these matched and unmatched states indicate the locations of meiotic crossovers.

**Figure 1.**
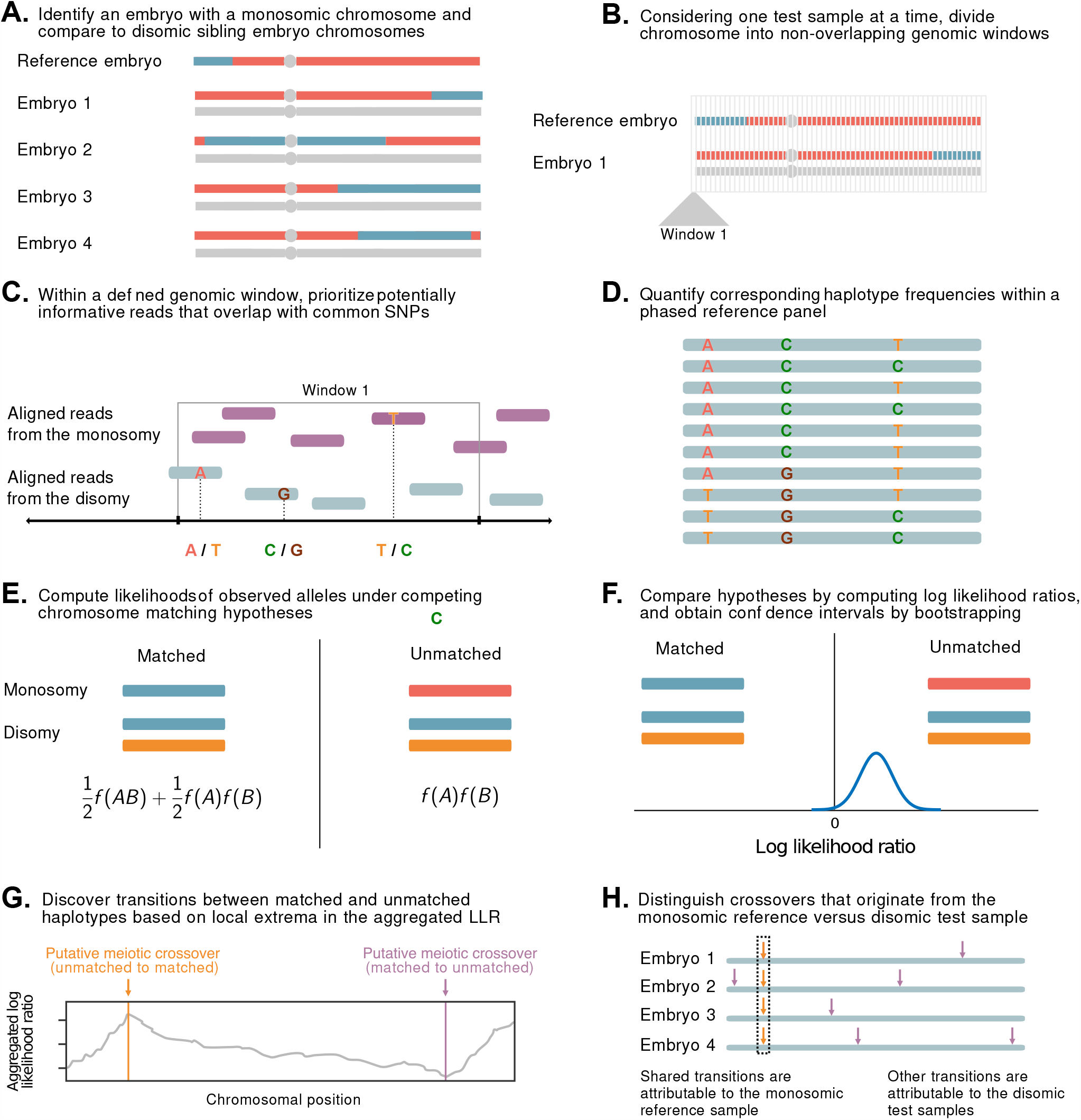
A statistical approach for meiotic crossover discovery based on low-coverage sequencing data from preimplantation genetic testing. A. Crossover detection is based on haplotype matching between a monosomic chromosome (which is phased by default) and the disomic chromosomes of sibling embryos from the same IVF case. B. Analysis is conducted within non-overlapping genomic windows on the scale of 10 - 100 kbp, defined by the length of typical human haplotypes. C. Within each window, 2-18 reads are re-sampled, prioritizing potentially informative reads that overlap common polymorphisms in the population. D. Frequencies and joint frequencies (i.e., haplotype frequencies) of these SNPs are quantified within an external phased genetic reference panel. E. Based on these frequencies, the likelihoods of the observed reads are computed under both the matched and unmatched haplotype hypotheses. F. The hypotheses are compared by computing a likelihood ratio, with variance estimated by bootstrapping. G. Local extrema in the aggregated log likelihood ratio indicate the locations of meiotic crossovers. H. Putative crossovers observed in the majority of sibling embryos can be attributed to the monosomic reference chromosome, while the remaining crossovers are attributed to the test samples.

### Evaluating method performance via simulation

In order to assess the performance of LD-CHASE, we simulated chromosomes from pairs of embryos consisting of a monosomy (i.e., reference sample) and disomy (i.e., test sample) that either shared a matching haplotype or were unrelated. A meiotic monosomy can occur due to errors at several distinct stages of oogenesis (Figure S2). We generated these pairs by mixing phased chromosomes from the 1000 Genomes Project [31], as described in the Methods. We focused our simulations on Chromosome 16, which is the chromosome most frequently affected by aneuploidy in preimplantation embryos, using a bin size of 2 Mbp and varying the sample ancestries across all superpopulations from the 1000 Genomes Project.

We allowed our classifier to assign bins as “matched”, “unmatched”, or “ambiguous” to denote uncertainty, and we used a balanced ROC curve to evaluate performance (see Methods; Figures 2 and S3). Our results demonstrated high sensitivity and specificity across all ancestries at a coverage of 0.05*×* per homolog (average area under the curve [AUC] of 0.989). As coverage was reduced to 0.025*×* and 0.013*×*, the AUC decreased by 0.014 and 0.053 on average, respectively, though performance was more or less affected in certain regions of the genome (Figure 2). Notably, performance of the classifier is affected by the local density of SNPs and sequencing reads, as well as ancestry matching between the reference panel and the target samples (Figures S3 and S4).

**Figure 2.**
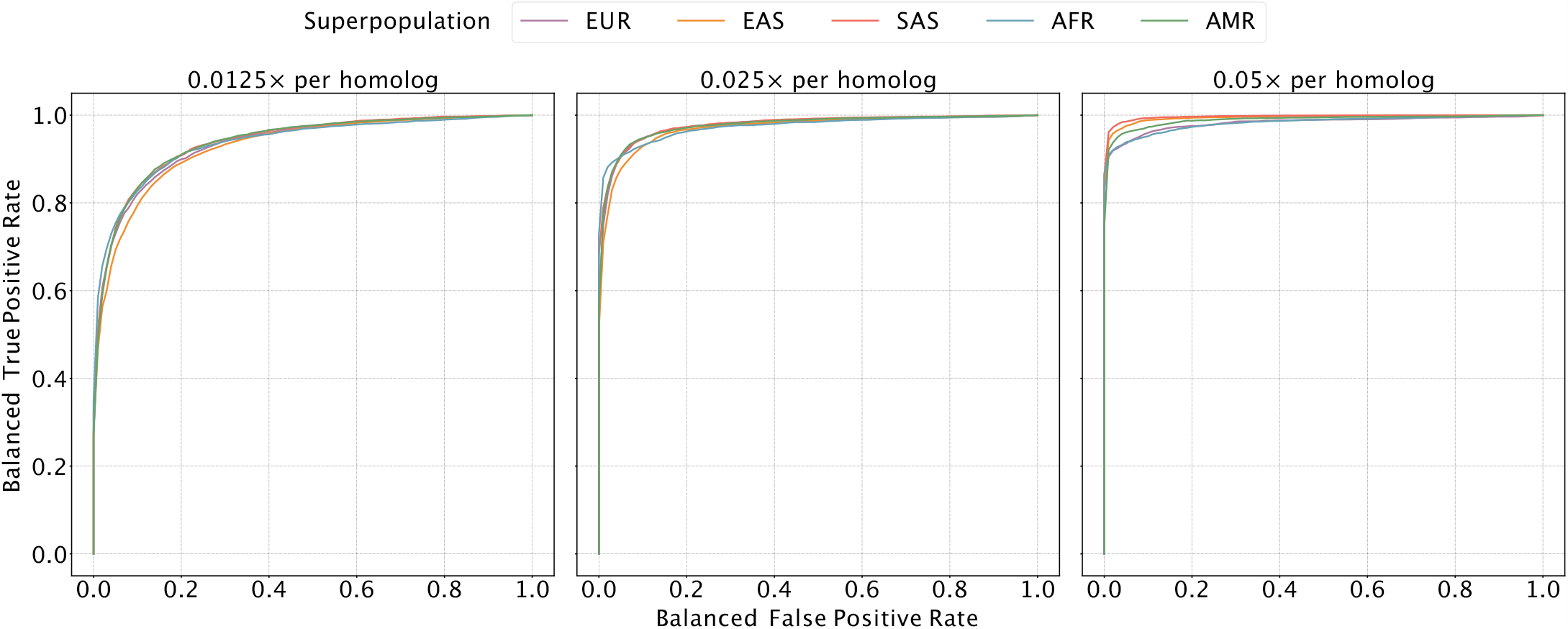
Evaluating the sensitivity and specificity of meiotic crossover detection based on simulation. Using data from the 1000 Genomes Project, we simulated pairs of Monosomy 16 and Disomy 16, where half of the pairs possessed matched haplotypes and the other half possessed unmatched haplotypes. Then we divided Chromosome 16 into 45 bins (of *∼* 2 Mbp) and calculated a balanced ROC curve (see Methods) for each bin, averaging over all the balanced ROC curves to obtain a mean balanced ROC curve. We repeated this procedure over a range of depths of coverage and across sets of samples from all superpopulations of the 1000 Genomes Project, abbreviated as follows: AMR = Admixed American; AFR = African; EAS = East Asian; EUR = European; SAS = South Asian.

### Application to a large PGT-A dataset

Encouraged by the performance of LD-CHASE on simulated data, we proceeded to apply it to a large dataset from the CReATe Fertility Centre (Toronto, Canada). The dataset consists of low-coverage sequencing data from 18,967 embryos from 2558 IVF patients, collected between April 2020 and August 2022. In order to select appropriate ancestry-matched reference panels, we first inferred the genetic similarity of each embryo to reference samples from the 1000 Genomes Project [31] using LASER [32] (see Methods). The results reflect the diverse ancestry composition of the patient population, with 68.71% (13,104) of embryos exhibiting the greatest genetic similarity to European reference samples, 6.76% (1290) of embryos exhibiting the greatest genetic similarity to South Asian reference samples, 5.74% (1094) of embryos exhibiting the greatest genetic similarity to East Asian reference samples, and the remaining embryos exhibiting lower genetic similarity to reference samples, for example due to recent admixture (Figure S5).

Previous studies have demonstrated that the vast majority of monosomies observed in blastocyst-stage embryos with PGT-A are of maternal meiotic origin, such that only the paternal chromosome remains [33, 21]. LD-CHASE uses such monosomies to map paternal crossovers in sibling disomic embryos. Meanwhile, haploidy or genome-wide uniparental isodisomy (GW-isoUPD) observed at the blastocyst stage nearly exclusively involves the sole presence of the maternal genome [33, 21, 34], allowing us to map genome-wide maternal crossovers in sibling disomic embryos. LD-CHASE thus requires preliminary analysis to identify chromosome abnormalities based on signatures of altered depth of coverage and/or genotype observations.

To this end, copy number of each autosome of each sample was inferred using WisecondorX [35] based on within-sample normalized depth of coverage. Across the entire dataset, we identified 388,366 disomies, 3307 trisomies, 4294 monosomies, 332 segmental gains, and 685 segmental losses (Figure S6 and Table S1). The monosomic chromosomes traced to embryos obtained from 1506 (58.85%) unique patients, facilitating mapping of paternal crossovers among 30,645 disomic chromosomes of 12,348 total embryos. Because maternal meiotic monosomies observed in blastocyst-stage embryos are highly enriched for Chromosomes 15, 16, 21, and 22, the mapping of paternal crossovers was largely relegated to these chromosomes, with much lower resolution for the rest of the genome.

Importantly, methods such as WisecondorX compare coverage across chromosomes within a sample and may therefore fail to detect aneuploidies that simultaneously affect many chromosomes. In extreme cases such as triploidy and haploidy/GW-isoUPD, where coverage is uniform across the genome despite the ploidy aberration, embryos may be erroneously classified as euploid. To overcome this limitation, we applied our published haplotype-aware method, LD-PGTA [28], to reclassify all chromosomes that were initially identified as disomic by WisecondorX. LD-PGTA identified 155 (1.65%) samples as triploid and 395 (4.20%) samples as haploid/GW-isoUPD. Importantly, such haploid/GW-isoUPD embryos were distributed across 184 (7.19%) patients, facilitating mapping of genome-wide maternal crossovers among 40,015 disomic chromosomes of 1898 total embryos.

### Sex-specific maps of crossovers on disomic chromosomes

Considering only chromosomes with informative genomic windows that covered at least 50% of their total length (see Methods), we identified 54,284 maternal crossovers across 27,026 chromosomes and 22,578 paternal crossovers across 21,050 chromosomes. An example of crossovers mapped in a single set of sibling embryos is provided in Figure 3, where transitions from intervals that do not match (blue) and match (red) the reference monosomic chromosome indicate the locations of meiotic crossovers (purple lines; see Methods). The exception to this interpretation involves transitions that are shared across all (or nearly all) sibling embryos, which instead reflect crossovers attributable to the reference monosomic chromosome itself (orange dashed lines). The genome-wide distributions of crossovers on disomic chromosomes are provided in Figures S7 and S8. We note that no crossovers are reported on the short arms of Chromosomes 13, 14, 15, 21, and 22 as their heterochromatic, highly repetitive nature makes them largely inaccessible to short read-based analyses. Moreover, for the same reason, these chromosome arms are largely devoid of variation in the reference panel data upon which our method relies.

**Figure 3.**
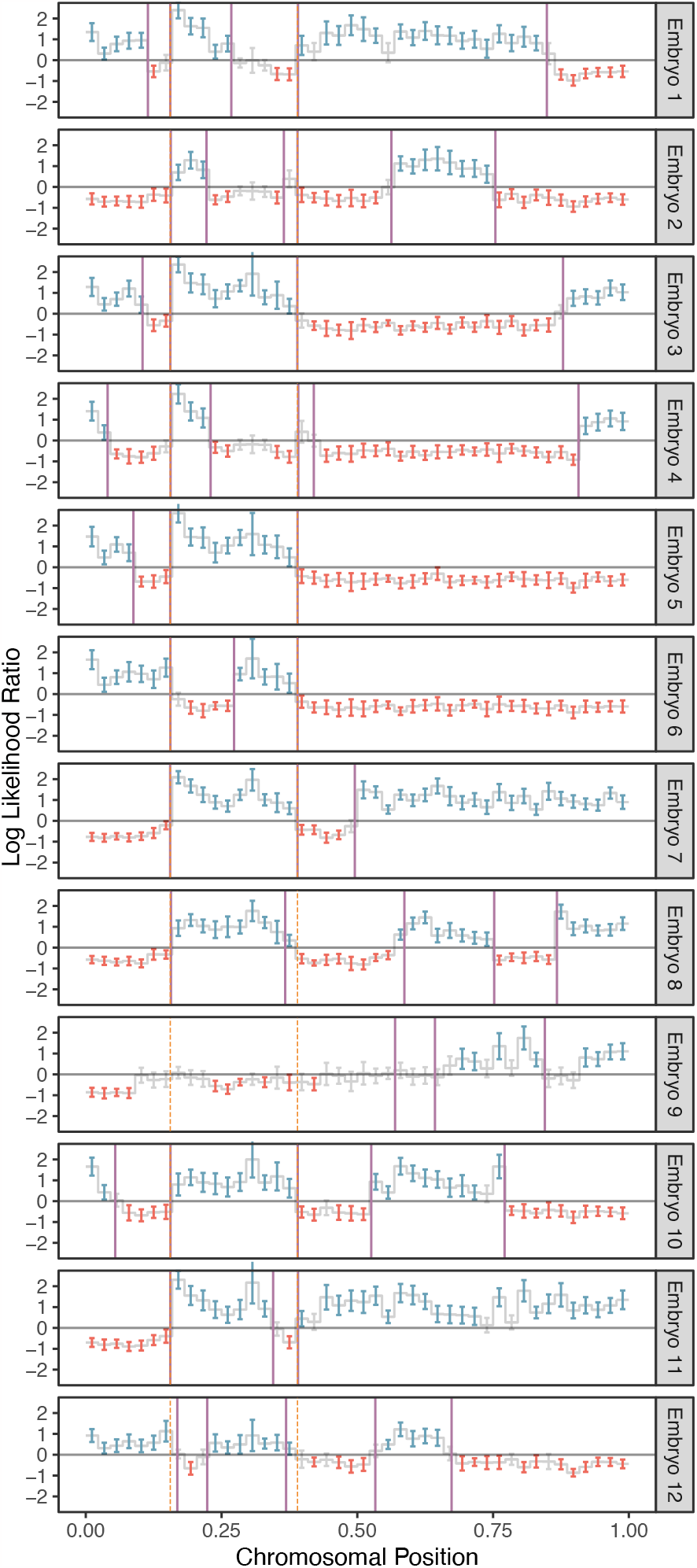
A representative example of crossover discovery based on haplotype matching among sibling IVF embryos. In this example, Chromosome 10 of each test embryo is compared to a single sibling reference embryo with monosomy of Chromosome 10. Evidence of haplotype non-matching is indicated by positive log-likelihood ratios, while evidence of haplotype matching is indicated by negative log-likelihood ratios. Crossovers are identified as transitions from positive (blue) to negative (red) log likelihood ratios or vice-versa. Each point corresponds to a bin size of 3 Mbp and consists of a varying number of genomic windows. Error bars denote 95% confidence intervals. Crossovers attributed to the monosomic chromosome are indicated with dashed orange lines (defined as those observed in more than half of test samples), while crossovers attributed to the test samples themselves are indicated with purple lines.

The crossover distributions were strongly correlated with sex-specific genetic maps published by deCODE, which were based on whole-genome sequencing of living parent-offspring trios [25], in broad support of the accuracy of our method (Figure 4). The observed correlation was particularly strong for putative maternal crossovers (*r* = 0.86) compared to putative paternal crossovers (*r* = 0.79). The smallest autosomal chromosomes are enriched for maternal meiotic aneuploidies and thus offer the greatest resolution for mapping paternal crossovers (see Figures S8 to S10 for chromosome-specific correlations). It was evident from our analysis that LD-CHASE performed differently for maternal and paternal crossovers because monosomies have only half the coverage compared to haploidy and uniparental isodisomy (isoUPD). This difference in depth of coverage stems from a lower copy number relative to the baseline. Additionally, the frequency of monosomies varies among autosomes, with some being rare. The low frequencies reduce our resolution and result in recombination maps that are relatively less precise. Notably, we observed that the correlations significantly declined when comparing our inferred female map to the deCODE male map (*r* = 0.33) and vice versa, supporting our assumptions about the parental origins of various chromosome abnormalities (Figures S11 and S12).

### Chromosome-specific propensities for various mechanisms of trisomy formation

Previous studies have suggested that chromosomes may vary in their susceptibility to segregation errors occurring during meiosis I (MI), meiosis II (MII), and mitosis [12, 13, 14, 36, 37, 38]. MI and MII errors can be roughly identified based on tracts of distinct (i.e., “both parental homologs” or BPH) or identical (i.e., “single parental homolog” or SPH) haplotypes, respectively, inherited from a single parent in regions spanning the centromere. Meanwhile, patterns of SPH chromosome-wide indicate a potential mitotic origin of trisomy (or MII error without recombination). While previous studies have noted that the attribution of centeromere-spanning BPH and SPH patterns to MI and MII errors is imperfect due to alternative mechanisms by which the signatures may originate [39], our results support the hypothesis that chromosomes possess unique propensities for various forms of segregation error (*x*^2^ [21, *N* = 1911] = 393.3, *P* = 2.3 *×* 10^*-*70^; Figure 5; Table S2). The vast majority (*>* 81%) of trisomies of Chromosomes 15, 16, 19, 21, and 22 exhibited haplotypic patterns consistent with errors in MI, while Chromosomes 11, 13, and 14 exhibited more modest excesses (*∼*70%) of MI errors (binomial test, Bonferroni adjusted-P *<* 0.05 for all noted chromosomes). Meanwhile, the remainder of chromosomes were characterized by a roughly equal number of MI and MII errors (36-61%; binomial test, Bonferroni adjusted-P *>* 0.05 for all noted chromosomes).

### An altered landscape of crossovers among aneuploid versus disomic chromosomes

Abnormal number or location of meiotic crossovers between homologous chromosomes may predispose oocytes to aneuploidy, as demonstrated by several previous studies [reviewed in 40]. Our published method, LD-PGTA [28], facilitates the mapping of crossovers on trisomic chromosomes, which can then be compared to the crossover map for disomic chromosomes obtained via LD-CHASE. One caveat of this comparison is that LD-PGTA and LD-CHASE possess different sensitivities and specificities, which also vary along the genome, as evident from our simulation-based benchmarking analyses (Figures S3, S4, S13 and S14). To ensure that observed differences between the crossover distributions were not driven by these technical differences, we merged together monosomies and disomies to create artificial trisomies (see Methods) such that the disomies and trisomies could both be analyzed with LD-PGTA in a standardized manner. The number (*r* = 0.99) and genome-wide distribution (*r* = 0.90) of disomic crossovers inferred by LD-CHASE and LD-PGTA exhibited strong agreement, suggesting that the methods are robust to their technical differences and supporting the use of LD-CHASE in downstream comparisons between trisomies and disomies (see Figure S15).

Across all chromosomes, these comparisons revealed a 35% depletion of crossovers for trisomies relative to disomies (Figure 6). On a per-chromosome basis, the depletion was observed across all chromosomes, but was largest for Chromosome 16 (54%) and smallest for Chromosome 4 (17%). While these observations are consistent with the hypothesis that a reduced rate (or absence; i.e., “exchangeless homologs”) of recombination contributes to aneuploidy risk [41, 40], we note that they may be partially driven by a failure of our method to detect crossovers on reciprocal recombinant chromosomes—a limitation that uniquely applies to trisomies but not disomies and affects nearly all previous genotype-based studies (see Discussion).

In addition to these global differences in numbers of crossovers, several chromosome-specific alterations in the landscapes of crossovers were evident from our results (Figure 6 and Figures S16 to S19). We used the Kolmogorov–Smirnov (KS) test to quantify differences in crossover landscapes, generating the null distribution by permutation (see Methods). Chromosomes 7, 14, and 16 exhibited significant differences in crossover landscapes between disomic and trisomic chromosomes (*p*-value *<* 0.05; though note small sample sizes for Chromosomes 7 and 14), whereas Chromosome 22 fell just above this threshold (*p*-value = 0.051; Table S3). Our observations reveal that on Chromosome 16, trisomies lacked a pair of hotspots in the vicinity of the centromere—one on each arm—broadly consistent with the previous observation that distal crossovers were enriched among a smaller sample of 62 cases of Trisomy 16 [13, 40]. Meanwhile, trisomies of Chromosome 22 appeared relatively enriched for crossovers near the center of the q-arm. Results for less frequent trisomies are depicted in Figures S16 to S19.

## Discussion

Meiotic crossovers are necessary for ensuring accurate pairing and subsequent segregation of chromosomes following decades-long dictyate arrest in human females [42]. Previous studies have demonstrated chromosome-specific associations between patterns of recombination and various forms of meiotic aneuploidy [37, 12, 13, 36, 38, 14]. For example, early studies of Trisomy 16 (the most common autosomal trisomy detected in human preimplantation embryos) suggested a depletion of recombination in pericentromeric regions relative to euploid control samples [43, 44]. However, the accuracy, genomic resolution, and statistical power of many such studies have been limited by the genetic assays they employed (e.g., Southern blot, PCR, etc.), as well as the challenge of achieving large samples from living trisomic individuals or products of conception. Due to its inherent relatedness structure, PGT-A offers a natural source of retrospective data for crossover mapping—including both viable and nonviable embryos—but current implementations based on low-coverage whole-genome sequencing pose a technical challenge for recovering relevant genotype information.

To address this challenge, our haplotype-aware method (LD-CHASE) uses known LD structure from an external reference panel as well as the frequent occurrence of monosomies (which are phased by default) to map crossovers based on comparisons of haplotypes among samples of sibling embryos. The resulting sex-specific maps of crossovers generated by our method were broadly consistent with those generated in previous prospective studies. The sex-specific nature of these patterns also supports our assumption that most monosomies observed in blastocyst-stage IVF embryos (of adequate morphology to be candidates for transfer and thus tested with PGT-A) originate during maternal meiosis, whereas most cases of haploidy/GW-isoUPD solely possess a maternally inherited set of chromosomes. The latter phenomenon may arise when sperm cells trigger egg activation but fail to fuse with the ovum, after which the maternal genome may duplicate to produce two identical complements [45]. Our observations about the sex-specific origins of various forms of aneuploidy thus independently replicate previous studies that directly assayed parental genomes [33, 21]. While we are eager to understand the differences in recombination maps of trisomies stemming from both MI and MII errors, the size of our dataset limits such exploration. The most frequently observed trisomies are 15, 16, 19, 21, and 22, yet fewer than 25% are of MII origin. In the future, we intend to delve deeper using larger PGT-A datasets that include parental genomic data. Furthermore, the ability to produce sex-specific crossover maps from PGT-A data could pave the way for studies seeking to understand the genetic basis of recombination phenotypes and the implications of extreme deviations in these phenotypes for meiotic errors and potential infertility.

As a first step toward this goal, we investigated the association between crossover phenotypes and chromosome abnormalities observed in preimplantation embryos. While broadly consistent with previous studies that used smaller samples or were restricted to living individuals with viable trisomies, our data provide unified views of these phenomena across all autosomes and support the hypothesis that the number and chromosome-specific locations of meiotic crossovers influence risk of aneuploidy. However, the observation that the length of the genetic map is shorter for trisomies versus disomies could be partially driven by the inability to detect crossovers on reciprocal chromosomes that derive from a single crossover event and were both transmitted to the oocyte to produce a trisomy. Future studies may employ linked-read or long-read sequencing to achieve direct read-based phasing and overcome this limitation [46, 47].

Despite the methodological advances we report here, crossover mapping from sequencing-based PGT-A data possesses several other technical limitations, including the modest resolution per sample (*∼*150 kbp). This limitation is driven by the combination of the low coverage (and thus sparsity) of the aligned reads (*<*0.05*×*), the low rates of heterozygosity of human genomes (*<*0.001), as well as the extent of LD in the reference panel (*<*1 Mb). Our benchmarking analyses additionally demonstrate that performance degrades with genetic distance between the test sample and reference panel due to differences in allele frequencies and LD structure, similar to challenges encountered when transferring polygenic scores across populations [48]. To overcome these sources of error, we aggregated signal across consecutive genomic windows, thereby increasing the classification accuracy at the cost of resolution.

Even with such an approach, it is important to note that performance is not uniform across the genome, as regions near centromeres, telomeres, or other repetitive regions are enriched for false positives and negatives relative to the genomic background.

An additional potential caveat regards the possibility that ovarian stimulation or other aspects of IVF may alter the crossover landscape of IVF embryos compared to non-IVF embryos or live-born individuals. While we cannot formally rule out this possibility, we consider it implausible due to the fact that maternal crossovers are established during prophase I of meiosis, which begins during female fetal development. As such, the locations of crossovers are determined long before any IVF-related procedures are introduced. Observed differences in the crossover landscapes between IVF embryos and living individuals are therefore likely to be indirect, for example driven by differences in viability of embryos with high versus low recombination rates due to the relationship between recombination and aneuploidy. While we expect this viability selection to apply to both IVF and naturally conceived embryos, this is an intriguing question for future investigation.

One promising future direction is the extension of our haplotype-aware approach to PGT-M, combining knowledge of population genetic patterns of LD in a reference panel with information from sibling embryos (or alternative data sources such as gametes) to infer transmission or non-transmission of pathogenic haplotypes from parents to offspring. While potentially lowering costs and increasing efficiency, such an approach will require extensive validation and benchmarking to determine its feasibility and accuracy given the probabilistic nature of LD and the high stakes of PGT-M. In the meantime, LD-CHASE offers a flexible tool for mapping crossovers in low-coverage sequencing data from multiple sibling embryos, toward a better understanding of the factors that modulate the meiotic crossover landscape and the role of recombination in the origins of aneuploidies.

## Methods

### Prioritizing informative reads

Our method seeks to overcome the sparse nature of low-coverage sequencing data by leveraging LD structure of an ancestry-matched reference panel, consisting of phased haplotypes from high-coverage sequencing data. Measurements of LD require pairwise and higher order comparisons and may thus grow intractable when applied to large genomic regions. To ensure computational efficiency, we developed a scoring algorithm to prioritize reads based on their potential information content, as determined by measuring haplotype diversity within a reference panel at sites that they overlap. We emphasize that the priority score of a read only depends on variation within the reference panel and not on the alleles that the read possesses. The score of a read is calculated as follows:

1. Given the ancestry composition of the target sample (e.g., 30% ancestry with genetic similarity to European reference samples and 70% ancestry with genetic similarity to East Asian reference samples), a suitable reference panel is chosen (see subsequent section titled “Assembling an ancestry-matched reference panel”).
2. Based on this reference panel, we list all biallelic SNPs that overlap with the read and their reference and alternative alleles.
3. Using the former list, we enumerate all the possible haplotypes. In a region that contains *n* biallelic SNPs, there are 2^*n*^ possible haplotypes.
4. The effective frequency of each haplotype is estimated from the reference panel:

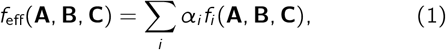

where *α*_*i*_ is the ancestry proportion from the *i* th population, and Σ_*i*_α_*i*_ =1 (e.g., *α*_1_ = 0.3 and *α*_2_ = 0.7). Moreover, *f*_*i*_ (**A, B, C**) is the joint frequency of the SNP alleles **A**,**B** and **C** in the *i* th population. Here we assumed *n* = 3, but the formula is applicable to any *n*.
5. We increment the priority score of a read by one for every haplotype with a frequency between *f*_0_ and 1 - *f*_0_.

An example of scoring a read that overlaps with three SNPs appears in the Supplemental Methods. Our scoring metric is based on the principle that reads that overlap SNPs with intermediate allele frequencies should receive high priority, as the inclusion of such sites will increase our ability to discern between two haplotypes. In the simplest case, where a read overlaps with only a single SNP, the score of the read would be two when the minor allele frequency (MAF) is at least *f*_0_ and otherwise zero. We note that all observed alleles from the same read are considered as originating from the same underlying molecule. Hence, our score metric reflects the number of common haplotypes existing in the population in the chromosomal region that overlaps with the read. For a reference panel on the scale of the 1000 Genomes Project (*∼*2500 unrelated individuals), 25% -45% of common SNPs have a nearest neighbor within 35 bp. Hence, even for short reads, it is beneficial to use a scoring metric that accounts for reads that span multiple SNPs.

### Comparing haplotype matching hypotheses

By virtue of LD, observations of a set of alleles from one read may provide information about the probabilities of allelic states in another read that originated from the same DNA molecule (i.e., chromosome). In contrast, when comparing reads originating from distinct homologous chromosomes, allelic states observed in one read will be uninformative of allelic states observed in the other read. As two siblings are characterized by chromosomal regions with different counts matched haplotypes, a change in the count along the chromosome indicates the position of a meiotic crossover, i.e., an exchange of DNA segments between non-sister chromatids during prophase I of meiosis. Similarly, a pair of half-siblings allows us to contrast the crossovers in either the paternal or the maternal homologs. Extending this logic, we note that sequences from either a monosomic sibling-embryo, a sibling embryo with GW-isoUPD or individual parental gametes would similarly allow us to isolate crossovers in embryo genomes that arose during oogensis or spermatogenesis.

We consider two sibling embryos, one has a monosomy and the second is a healthy diploid, consisting of two copies of the genome—a maternal and a paternal copy. For a set of reads aligned to a defined genomic region, we compare the likelihoods of the observed alleles under two competing hypotheses:

1. The monosomic embryo and the healthy diploid have matched haplotypes, denoted as matched haplotypes hypothesis.
2. The monosomic embryo and the healthy diploid have unmatched haplotypes, denoted as unmatched haplotypes hypothesis.

A transition between regions with matched and unmatched haplotypes indicates the location of a meiotic crossover.

Our statistical models consider a situation where one read is drawn from the monosomic chromosome and the second from the disomic chromosome of a sibling. The odds of two reads being drawn from identical versus distinct haplotypes differ under the matched and unmatched haplotypes hypotheses. Specifically, for the matched haplotypes hypothesis, the odds are 1: 1, and for the unmatched haplotypes hypothesis, the odds are 0: 1. If a pair of reads is drawn from identical haplotypes, the probability of observing the two alleles is given by the joint frequency of these two alleles (i.e., the frequency of the haplotype that they define) in the reference panel. In contrast, if a pair of reads is drawn from distinct haplotypes, then the probability of observing the two alleles is simply the product of the frequencies of the two alleles in the reference panel:

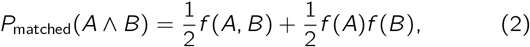

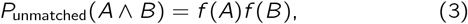

where *f* (*A*) is the frequency of allele *A* and *f* (*A, B*) is the joint frequency of alleles *A* and *B* in the population.

These statistical models can then be generalized to arbitrary admixture scenarios by a simple substitution. We assume that each distinct parental haplotype is drawn from an ancestral population with a probability equal to the ancestry proportion of the tested individual that is associated with that population. In accordance with the assumption, we replace each allele frequency distribution, *f* by the combination Σ_*i*_ *α*_*i*_ *f*_*i*_. Here *α*_*i*_ is the probability that the alleles originated from the *i* -th population, *f*_*i*_ is the allele frequency distribution for the *i* -th population and Σ_*i*_ *α*_*i*_ = 1. For example, under this substitution *P*_unmatched_(*A ^ B*)= *α*_1_*f*_1_(*A, B*)+ (1 − *α*_1_)*f*_2_(*A, B*), for admixture between two populations.

Likelihoods of the two hypotheses are compared by computing a log-likelihood ratio:

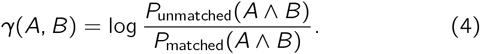

When a read overlaps with multiple SNPs, *f* (*A*) should be interpreted as the joint frequency of all SNP alleles that occur in read *A* (i.e., the frequency of haplotype *A*). Similarly, *f* (*A, B*) would denote the joint frequency of all SNP alleles occurring in reads *A* and *B*. The equations above were extended to consider up 6 reads per window and homolog, as described in the later subsection titled “Generalization to arbitrary number of reads”. Estimates of allele and haplotype frequencies from a reference panel do not depend on theoretical assumptions, but rely on the idea that the sample is randomly drawn from a population with similar ancestry. One limitation, which we consider, is that reliable estimates of probabilities near zero or one require large reference panels, such as the 1000 Genomes Project [31].

### Determining optimal size of a genomic window

Because pairwise LD in human genomes decays on average to a quarter of its maximal value over physical distances of 100 kbp [31], the length of the chromosomes is divided into genomic windows on a scale consistent with the length of typical human haplotypes (10^4^ - 10^5^ bp). While one library consists of two homologs and the second consists of a single homolog, we would like to sample an even number of reads from each homolog. Thus, for each DNA sequence, we only consider the depth of coverage per homolog (i.e., we divide the coverage by the ploidy for the chromosome of each sample under consideration).

We require a minimal number of reads per genomic window and homolog, as determined by the sample with the lowest average depth of coverage. We then scan the chromosome in a sliding window, using a window size that adjusts according to the local depth of coverage of the two different sequenced samples. This adaptive sliding window approach possesses advantages over a fixed length window in that it (a) accounts for GC-poor and GC-rich regions of a genome, which tend to be sequenced at lower depths of coverage using Illumina platforms [49] and (b) accounts for varying densities of SNPs across the genome [50].

The algorithm simultaneously scans aligned reads from the two samples in the forward direction of the reference genome and identifies informative reads (i.e., reads that reach the priority score threshold) from both samples within genomic windows. For each genomic window, the minimal number of required reads from the DNA sequence of the disomy is twice the minimal number from the monosomy. If (a) the distance between consecutive reads in one of the samples exceeds 100 kbp or (b) a genomic window extends to 350 kbp and does not meet the minimal number of reads per homolog, the window is dismissed.

### Quantifying uncertainty by bootstrapping

To quantify uncertainty in our likelihood estimates, we performed *m* out of *n* bootstrapping by iteratively resampling reads within each window [51]. Resampling was performed without replacement to comply with the assumptions of the statistical models about the odds of drawing two reads from the same haplotype. Thus, in each iteration, only subsets of the available reads can be resampled. Specifically, within each genomic windows, up to 6 reads per homolog with a priority score exceeding a defined threshold are randomly sampled with equal probabilities. The likelihood of the observed combination of SNP alleles under each competing hypothesis is then calculated, and the hypotheses are compared by computing a log-likelihood ratio (LLR). The sample mean and the unbiased sample variance (i.e., with Bessel’s correction) of the LLR in each window are calculated by repeating this process using a bootstrapping approach,

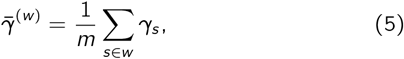

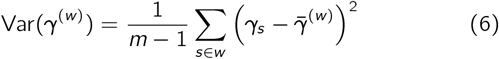

where, *y*_*s*_ is the the log-likelihood ratio for *s*-th subsample of reads from the *w*-th genomic window and *m* is the number of subsamples. Because the number of terms in the statistical models grows exponentially, we subsample at most 6 reads per window and homolog. Moreover, accurate estimates of joint frequencies of many alleles requires a very large reference panel. Given the rate of heterozygosity in human populations and the size of the 1000 Genomes Project dataset, 6 reads per homolog is generally sufficient to capture one or more heterozygous SNPs that would inform our comparison of hypotheses.

### Aggregating signal across consecutive windows

Even when sequences are generated according to one of the hypotheses, a fraction of genomic windows will emit alleles that do not support that hypothesis and may even provide modest support for an alternative hypothesis. This phenomenon is largely driven by the sparsity of the data, as well as the low rates of heterozygosity in human genomes, which together contribute to random noise. Another possible source of error is a local mismatch between the ancestry of the reference panel and the tested sequence. Moreover, technical errors such as spurious alignment and genotyping could contribute to poor results within certain genomic regions (e.g., near the centromeres). To overcome this noise, we binned LLRs across consecutive genomic windows, thereby reducing biases and increasing the classification accuracy at the cost of resolution. Specifically, we aggregated the mean LLRs of genomic windows within a larger bin,

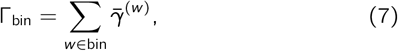

where 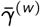 is the mean of the LLRs associated with the *w*-th genomic window. In addition, using the Bienaymé formula, we calculated the variance of the aggregated LLRs,

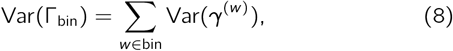

where Var(*γ*^(*w*)^) is the variance of the LLRs associated with the *w*-th window. For a sufficiently large bin, the confidence interval for the aggregated LLR is 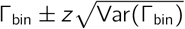, where *z* = Φ^−1^ (1 − *α*) is the *z* -score, <t is the cumulative distribution function of the standard normal distribution and *C* = 100(1 − 2*α*)% is the confidence level. The confidence level is chosen based on the desired sensitivity vs. specificity. We normalized the aggregated LLRs by the number of genomic windows that compose each bin, 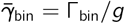. Thus, the variance of the mean LLR per window is Var 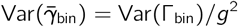. These normalized quantities can be compared across different regions of the genome, as long as the size of the genomic window is the same on average.

### Simulating parental haplotypes and diploid offspring

Using a generative model, we simulated trios of a (a) parental chromosome, (b) diploid offspring with haplotypes matching the parental chromosome and (c) an unrelated chromosome. This allows us to evaluate the classifier performance in each genomic window along the human genome. To this end, we constructed synthetic samples comprising combinations of phased haplotypes from the 1000 Genomes Project [31]. These phased haplotypes are extracted from variant call sets to effectively form a pool of haploid sequences.

We first consider non-admixed offspring by drawing 3 effective haploid sequences from the same superpopulation. The first two haploid sequences are used to simulate the diploid offspring, while the first and third sequences are used to simulate the parental chromosome and the unrelated chromosome, respectively. We then simulate reads by selecting a random position along the chromosome from a uniform distribution to represent the midpoint of an aligned read with a given length. Based on the selected position, one out of the three haplotypes is drawn from a discrete distribution,

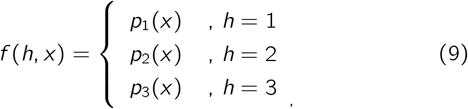

where, in general, the probability of haplotype *h* depends on the position of the read, *x*. When simulating a non-admixed diploid offspring, the first haplotype is just as likely as the second haplotype (*p*_2_ = *p*_1_), and the third haplotype is absent (*p*_3_ = 0). Similarly, for the parental chromosome, *p*_1_ =1 and *p*_2_ = *p*_3_ = 0, while for the unrelated chromosome, *p*_3_ =1 and *p*_1_ = *p*_2_ = 0. Then, from the selected haplotype, *h*, a segment of length *l* that is centered at the selected chromosomal position, *x*, is added to simulated data, mimicking the process of short-read sequencing. This process of simulating sequencing data is repeated until the desired depth of coverage is attained.

In order to simulate an offspring descended from parents from distinct superpopulations (hereafter termed “recent admixed ancestry”), we draw two effective haploid sequences from different superpopulations of the 1000 Genomes Project. A third haploid sequence is then drawn from one of the two former pools to simulate the unrelated chromosome. Finally, we use the generative model with these three effective haploid sequences to simulate reads. A procedure for simulating diploid offspring under a scenario involving more distant admixture is discussed in the Supplemental Methods.

### Evaluating model performance on simulated data

We developed a classification scheme to determine whether a bin supports one of two competing hypotheses. To this end, we performed *m* out of *n* bootstrapping by iteratively resampling reads within each window pairs and computed log likelihood ratios (LLRs) of competing statistical models, as described in the subsection titled “Quantifying uncertainty by bootstrapping”.

The confidence interval for the mean LLR is 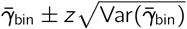, and *z* is referred to as the *z*-score. Thus, we classify a bin as exhibiting support for the matched haplotypes hypothesis when

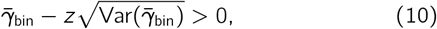

and for the unmatched haplotypes hypothesis when

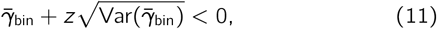

where the first (second) criterion is equivalent to requiring that the bounds of the confidence interval lie on the positive (negative) side of the number line. When a confidence interval crosses the origin of the number line, we classify the bin as ambiguous (see Figure S20 for a diagram of these classes).

For a given depth of coverage and read length, we simulate an equal number of sequences generated according to both hypotheses, as explained in the previous subsection. We define true positives (negatives) as simulations where sequences generated under the “unmatched” (“matched”) haplotypes hypothesis are correctly classified. Based on these simulations, we generate balanced receiver operating characteristic (ROC) curves for each bin [28]. The balanced true and false positive rates for a bin are defined as

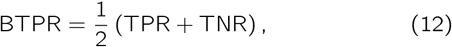

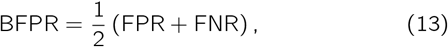

where TPR, TNR, FPR and FNR the true positive rate, true negative rate, false positive rate and false negative rate, respectively.

The “balanced ROC curve” is tailored for trinomial classification tasks. Here the three possible classes are “unmatched”, “matched” and “ambiguous”. The ambiguous class contains all instances that do not fulfill the criteria in Equations (10) and (11) (i.e., instances where the boundaries of the confidence interval span zero). This classification scheme allows us to optimize the classification of both “unmatched” and “matched” instances at the expense of leaving ambiguous instances. The advantage of this optimization is a reduction in the rate of spurious classification. To generate a balanced ROC curve for each genomic bin, we varied the *z*-score.

### Generalization to an arbitrary number of reads

The model of the unmatched haplotypes hypothesis for non-admixed samples and an arbitrary number of reads is based on the disomy model for non-admixed samples that was derived for LD-PGTA [28]. More specifically, the “umatched” statistical model for *m* + *n* reads is merely the joint frequency *f* (*A*_1_, *A*_2_, …, *A*_*m*_) multiplied by the disomy model *P*_disomy_(*B*_1_ ^ *B*_2_ *^* …, *B*_*n*_) of *n*-reads for non-admixed samples. Here *A*_*i*_ are reads drawn from the monosomic reference sample, while *B*_*i*_ are reads drawn from the disomic test sample.

The model of the matched haplotypes hypothesis for non-admixed samples and an arbitrary number of reads is based on the disomy model for recent-admixtures, which was previously introduced in Ariad et al. [28]. More specifically, modeling the matched haplotypes hypothesis for *m* + *n* reads can be accomplished by substituting an effective joint frequency distributions in the disomy model of *n*-reads for recent-admixture. The adjusted model involves two distributions *f* (*X*) and *g*(*X*). The distribution *f* is derived from a reference panel of a population as before, while *g* is an effective distribution that is defined as *g*(*X*) *≡ f* (*A*_1_, *A*_2_,…, *A*_*m*_,*X*). The reads *A*_*i*_ with *i* = 1, 2,…,*m* are drawn only from the monosomic reference sample, while the rest of the reads are drawn from the disomic test sample. Also, each term in the linear model should take into account the presence of the reads *A*_1_, *A*_2_,…, *A*_*m*_ and thus terms in the disomy model for recent admixtures that involve only the distribution *f* should multiplied by *g*(?) *≡ f* (*A*_1_,…, *A*_*m*_). Derivations and explicit statistical models for non-admixed ancestry, recent admixture and more distant admixture scenarios for *m* + 2, *m* +3 and *m* +4 reads can be found in the Supplemental Methods.

### Identifying meiotic crossovers

In order to identify the locations of meiotic crossovers, we analyze the cumulative sums of log-likelihoods ratios (LLRs) from individual genomic windows as we move along the chromosome. Since we performed *m* out of *n* bootstrapping by iteratively resampling reads within each window and calculating LLRs, we define two quantities:

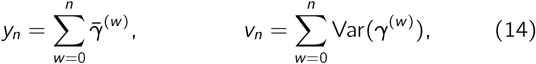

where we assume negligible linkage disequilibrium between alleles in different genomic windows, and hence according to the Bienaymé formula 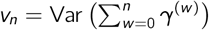. Local maxima of *y*_*n*_ indicate potential transitions from “unmatched” to “matched” regions, while local minima of *y*_*n*_ indicate potential transitions from “matched” to “unmatched” regions. Thus, a crossover occurred within the *j* th genomic window if there exists a region in which either arg max (*y*_*n*_)= *j* and in addition

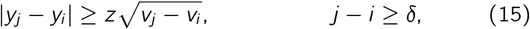

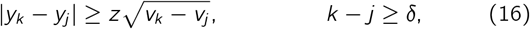

where *z* is the threshold for calling a crossover and *c5* is the minimal number of genomic window in the region. In addition, we define a metric that describes the confidence in calling the *j* th crossover:

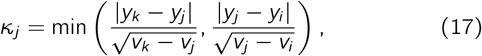

which fulfills *κ*_*j*_ *> z*_score_. When, based on the criteria above, two consecutive crossovers are identified as local maxima or minima of the accumulated LLR, *y*_*n*_, this implies that a crossover was skipped. Failure to detect a crossover occurs when the values of *z* and/or *c5* are too restrictive. Given two consecutive maxima in the *i* th and *k* th genomic windows, we identify the genomic window with skipped minimum as

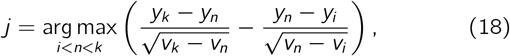

where this condition means that *z*_score_ of the “matched” and “unmatched” intervals are maximized simultaneously. In case of two consecutive minima, we replace the function arg max with arg min.

### Assigning crossovers to reference versus test samples

Because we scan for crossovers by comparing sequences from two sibling embryos, assigning each crossover to a single embryo requires additional information. When sequencing data from three or more sibling embryos is available, we contrast the crossovers of sibling embryos with one common monosomic sibling (see Figure 1). This, in turn, produces a repeated pattern in each sequence of crossovers that can be attributed to the common sibling (see Figure 3). Once the repeated pattern is identified, we subtract it to recover the crossovers in the rest of the embryos.

More specifically, we consider a set of *n* +1 sibling embryos, where one embryo is used to contrast the crossovers in the rest of the sibling embryos. All crossovers are then combined to form a sorted list, and we scan the list for *n* sequential crossovers within a region of size *l*. For each cluster, we calculate the average position of the crossovers. The average position from each cluster is associated with the common sibling. Finally, the union of all the clusters is subtracted from each of the *n* sequences of crossovers, and remaining crossovers are traced back to the sibling embryos.

The sequencing quality may vary from one embryo to another, and some crossovers might not be identified. Hence, we adjust the algorithm to allow clusters with various sizes. After seeking all clusters of size *n*, we continue seeking clusters of size *n-* 1. This process of seeking smaller clusters is repeated iteratively for cluster sizes greater than *n/*2. Another issue that may arise is that when the regions size, *l*, is too large, two or more crossovers from the same embryo may overlap. In such cases, we only consider the crossover that is closest to the cluster mean.

Each crossover that is attributed to the common reference embryo is assigned two scores. The first score is the proportion of sibling embryos supporting the crossover: )λ_*i*_ = *k*_*i*_ */n*, where *k*_*i*_ and *n* are the size of cluster *i* and number of contrasted embryos, respectively. The second score is the minimal confidence score in the *i* th cluster: 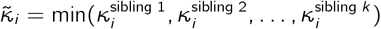, where κ was defined in the subsection titled “Identifying chromosomal crossovers”.

### Isolating sex-specific meiotic crossovers

Here we discuss the possibility of identifying sex-specific crossovers by matching haplotypes between natural occurring monosomies and genome-wide isodisomies (GW-isoUPD) in IVF embryos and disomic sibling embryos from the same IVF cycles. This in turn allows us to leverage the high volume of aneuploidies observed in IVF to obtain sex-specific distributions of crossovers.

We also review how maternal crossovers in trisomies can be identified by scanning for transitions between tracts of two versus three unique homologs.

Paternal crossovers (i.e., crossovers originating from spermatogenesis) can be identified by focusing on IVF cycles that yielded at least one embryo with a monosomy of a given chromosome, as well as at least one embryo that is disomic for that chromosome. The vast majority of monosomies are of maternal meiotic origin, such that the remaining chromosome is of paternal origin. The opposite scenario, whereby a haploid ovum is fertilized by an nullisomic sperm cell accounts for less than 10% of autosomal monosomies [5, 52, 53]. Moreover, when intracytoplasmic sperm injection (ICSI) is performed, sperm *in vitro* selection and capacitation may further reduce the rates of paternal-origin monosomies [54, 55, 56, 57]. Conversely, maternal crossovers (i.e., crossovers originating from oogenesis) can be identified by focusing on IVF cycles that yielded at least one case of haploidy/genome-wide isodisomy (which are indistinguishable with sequencing-based PGT-A), as well as at least one sibling that is disomic for one or more chromosomes. Such cases of GW-isoUPD may originate from oocytes that commence cleavage and early embryonic development without fertilization [34].

Aneuploidy (gain or loss of entire chromosomes) is the leading cause of IVF failure. We found that 351 (13.7%), 529 (20.7%), 332 (13.0%), and 470 (18.4%) of IVF cycles in the CReATe dataset had at least one embryo with monosomy of Chromosome 15, 16, 21, or 22, respectively, as well as 7.3 disomic siblings on average (see Table S4 for the rest of the autosomes). Thus, LD-CHASE can use these natural occurring monosomies to identify paternal crossovers along the chromosomes of sibling embryos. Similarly, we found that 201 (9.1%) of the IVF cycles in the CReATe dataset had at least one haploid/GW-isoUPD embryo and an additional 7.8 euploid siblings on average (see Table S5), offering good resolution for mapping maternal crossovers genome-wide.

### Coverage-based discovery of chromosome abnormalities

We used WisecondorX to deduce the chromosomal copy numbers of each of 20,160 embryos in the CReATe dataset [35, 58]. To this end, we first created four sets of reference samples for read counts: (a) 9202 sequences were obtained from biopsies of 4-6 cells and consist of single reads of 75 bp. (b) 9818 sequences were obtained from biopsies of 4-6 cells and consist of paired-end reads of 75 bp (c) 578 sequences were obtained from biopsies of 4-5 cells and consist of single reads of 75 bp, and (d) 562 sequences were obtained from biopsies of 4-5 cells and consist of paired-end reads of 75 bp. We used all the sequences in a category (e.g., sequences associated with 4-5 cells and single reads) as reference samples. This approach is effective as long as aneuploid chromosomes are rare and as long as the rate of chromosome loss and chromosome gain are sufficiently similar to balance out one another in large samples. The first assumption is justified based on previous PGT-A studies, which showed that the rate of aneuploidy per chromosome is less than 10% (including monosomy, trisomy and mosaics)[28]. The second assumption is justified by noting that for each chromosome, the rate of trisomy should be similar to the rate of monosomy, as both are mainly caused nondisjunction [5]. To assess the robustness of this approach, we compared the results obtained from WisecondorX with those generated by NxClinical, a diagnostic tool utilized by the CReATe Fertility Centre, for select chromosomes that exhibited copy number variations, as shown in Figure Figure S21. In addition, we applied WisecondorX to a separate published PGT-A dataset from the Zouves Fertility Center consisting 8881 samples. The dataset was previously analyzed using BlueFuse Multi, and we found that the inferred copy numbers were in strong agreement with the WisecondorX. The number of sequences that were analyzed successfully via WisecondorX was 20,114. The number of relevant sequences was further reduced to 18,967 after filtering sequences without a corresponding record in the metadata table or when the genetic ancestry could not be inferred.

### Haplotype-aware discovery of ploidy abnormalities

Coverage-based approaches for inferring chromosome copy numbers, such as WisecondorX, are based on relative differences in the depth of coverage across chromosomes within samples. As such approaches assume that the baseline coverage corresponds to disomy, scenarios when many chromosomes are simultaneously affected, such as haploidy and triploidy, violate this assumption and may elude detection. To overcome this limitation, we applied our haplotype-aware method, LD-PGTA, to scan for the number of unique haplotypes along the genome [28] and calculate chromosome-wide log-likelihood ratios (LLR) comparing hypotheses of monosomy, disomy, and various forms of trisomy. When the LLR for at least 15 chromosomes supported a common aneuploidy hypothesis (monosomy or trisomy), the sample was classified as haploid/GW-isoUPD or triploid, respectively.

### Assembling an ancestry-matched reference panel

Given the aforementioned importance of the ancestry of the reference panel, we used LASER v2.04 [32, 59] to perform automated ancestry inference for each embryo sample from the low-coverage sequencing data. LASER applies principal components analysis (PCA) to genotypes of reference individuals with known ancestry. It then projects target samples onto the reference PCA space, using a Procrustes analysis to overcome the sparse nature of the data. We excluded markers with less than 0.01*×* depth of coverage and restricted analysis to the top 32 principal components, performing 5 replicate runs per sample.

Ancestry of each target sample was deduced using a *k*-nearest neighbors approach. Specifically, we identified the nearest 150 nearest neighbor reference sample to each target sample based on rectilinear distance. We then calculated the superpopulation ancestry composition of the 150 reference samples. When >95% of such samples derived from the same 1000 Genomes Project superpopulation, we employed downstream statistical models designed for non-admixed samples. In cases where two superpopulations were represented in roughly equal proportions (maximal difference of 10%), we employed downstream statistical models designed for recent admixture. For all other samples, we employed downstream statistical models designed for more distant admixture scenarios. For this latter group, superpopulations represented at levels of ≥5% among the nearest neighbors to each target sample were used in the construction of reference panels.

### Testing robustness versus LD-PGTA

One aspect of our study involved comparing the landscapes of meiotic crossovers between disomic and trisomic chromosomes, as inferred by LD-CHASE and LD-PGTA, respectively. This comparison poses a statistical challenge, as the two methods possess distinct sensitivities and specificities, which also vary across the genome and as a function of coverage of the respective samples. To validate our findings, we therefore sought to compare these landscapes under a single statistical framework by combining disomic samples with separate monosomic samples to produce artificial trisomies. Such artificial trisomies can be analyzed uniformly with LD-PGTA and compared to true trisomies analyzed with the same method, as well as to the more direct analysis of disomies with LD-CHASE.

In principle, such artificial trisomies can be produced by combining monosomies and disomies from the same IVF patients in ratios of 2: 1, such that they can be analyzed with LD-PGTA. In practice, naive merging based on genome-wide average depth of coverage yields poor results due to the low complexity of DNA libraries from PGT-A, which results in sample-specific non-uniformity in coverage across the genome. To overcome this challenge, we merely replaced the statistical models of LD-CHASE with those of LD-PGTA, thereby controlling the ratio of reads from each sample on a local (i.e., genomic window) as opposed to genome-wide scale. Thus, the construction of the genomic windows and the approach to sample reads from a genomic window is identical to LD-CHASE. However, the statistical models of LD-PGTA have no prior knowledge about the DNA sample from which a given read is drawn. Figures S16 to S19 displays the concordance in crossover distributions between disomic chromosomes as analyzed by both LD-CHASE and LD-PGTA.

### The coverage of genomic windows

To further address the low complexity of DNA libraries prepared from few input cells, we introduced an additional metric that simultaneously captures both the depth of coverage and the complexity of the library. After we tile a chromosome with genomic windows, as described in “Determining optimal size of a genomic window”, we calculate the coverage of genomic windows for a given sample as:

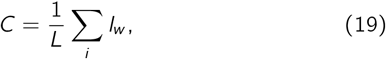

where *l*_*w*_ is the length of the *w* th genomic window, *L* is the length of the chromosome and thus 0 *<C <* 1. We then restricted our analysis to samples with *C* ≥ 0.5 (i.e., genomic windows covering at least half of the chromosome). For the subset of patients with multiple monosomic embryo samples (affecting the same chromosome) from which to choose, we selected the monosomy that yielded the highest value of *C* in order to maximize resolution for identifying meiotic crossovers.

### Distinguishing trisomies originating from errors in meiosis I and meiosis II

Samples with tracts of BPH in regions surrounding the centromere were classified as putative meiosis I errors, while tracts of BPH elsewhere on the chromosome were classified as putative meiosis II errors. Ambiguous samples were also noted. Specifically, regions bounded by crossovers and emitting a *z*-score above 1.96 and below *-*1.96 were regarded as BPH and SPH regions, respectively, and otherwise regarded as ambiguous. We defined a pericentromeric region as a region 20% of the chromosome length and centered on the centromere. However, for the acrocentric chromosomes, the pericentromeric region only includes the q-arm and is thus effectively reduced to 10% of the chromosome length. When at least 10% of the chromosome exhibited tracts of BPH, the trisomy was classified as a meiotic error. If it was not classified as a meiotic error and at least 50% of the chromosome exhibited tracts of SPH, the trisomy was classified as a mitotic error. Otherwise, it was classified as ambiguous. Cases that were classified as meiotic errors were further classified as follows: When at least 50% of the pericentromeric region reflected BPH, the trisomy was classified as a MI error. Moreover, if it was not classified as MI and at least 50% of the pericentromeric region reflected SPH, then the trisomy was classified as an MII error. Otherwise the case was classified as ambiguous. In this analysis, only trisomy cases with genomic windows that covered at least 50% of the chromosome length were taken into account.

To compare the frequency of MI versus MII error across chromosomes, we fit a binomial generalized linear model implemented with the ‘lme4’ package [60], where the response variable was defined as the counts of MI and MII errors per patient, patient identifier was included as a random effect predictor variable (to account for non-independence among sibling embryos), and chromosome was included as a categorical predictor variable. We compared this full model to a reduced model without the chromosome predictor variable using analysis of deviance.

### Comparing distributions of chromosomal crossover via eCDFs and KS test

Each chromosome in a given sample typically exhibits between 1 and 3 crossover events. Although the overall pool of crossovers exhibits a continuous spatial distribution along the genome, the underlying crossovers are largely independent events and thus can be treated as such in downstream statistical tests. To formulate our comparison of landscapes, we summarized each landscape as an empirical cumulative distribution function (eCDF), which traces the cumulative genetic map length as one moves from the beginning to the end of a given chromosome (i.e., a line plot comparing the physical map to the genetic map). One advantage of this approach is that it circumvents the need to bin the data and thus is bin-size independent. We note that such summaries are common in the recombination literature, e.g., Fig. 4b of Peñalba and Wolf [2]. We then applied the two-sample Kolmogorov–Smirnov (KS) test to test whether two underlying one-dimensional probability distributions differ, computing the *p*-value by permutation. While the KS test is nonparametric and makes no assumptions about the form of the distribution from which the data were drawn, permutations further ensure that the *p*-value calculation is based solely on the observed data without any reliance on asymptotic approximations. To calculate the *p*-value, we constructed a null hypothesis that posits the two empirical samples come from the same continuous distribution. This was achieved by evaluating all possible combinations of assignments of the combined data into two groups, each of the sizes of the two original samples, and computing the KS statistic for each combination. The *p*-value is then computed as the proportion of permutations that result in a KS statistic as extreme as, or more extreme than, the observed statistic.

**Figure 4.**
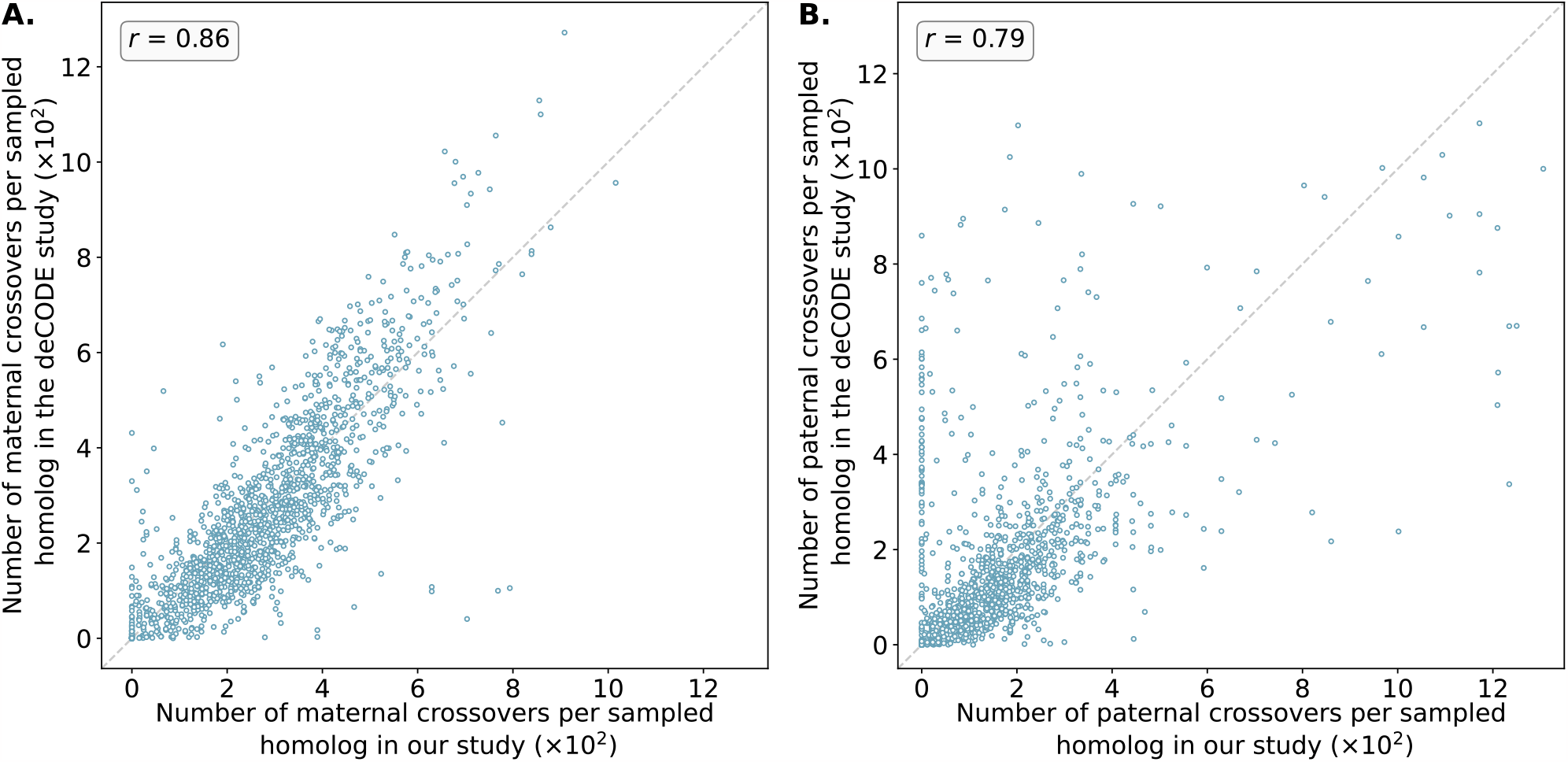
Number of maternal (A) and paternal (B) crossovers per sampled homolog in genomic bins compared between our study and published genetic maps from deCODE. Data from deCODE were obtained from Halldorsson et al. [25]. Crossovers were identified as transitions between regions of “matched” and “unmatched” haplotypes, where each region included at least 15 genomic windows and a *z*-score of the at least 1.96. Pearson correlation coefficients (*r*) between the deCODE map and our map across genomic bins are reported for each panel (See Figures S9 and S10 for chromosome-specific comparison).

**Figure 5.**
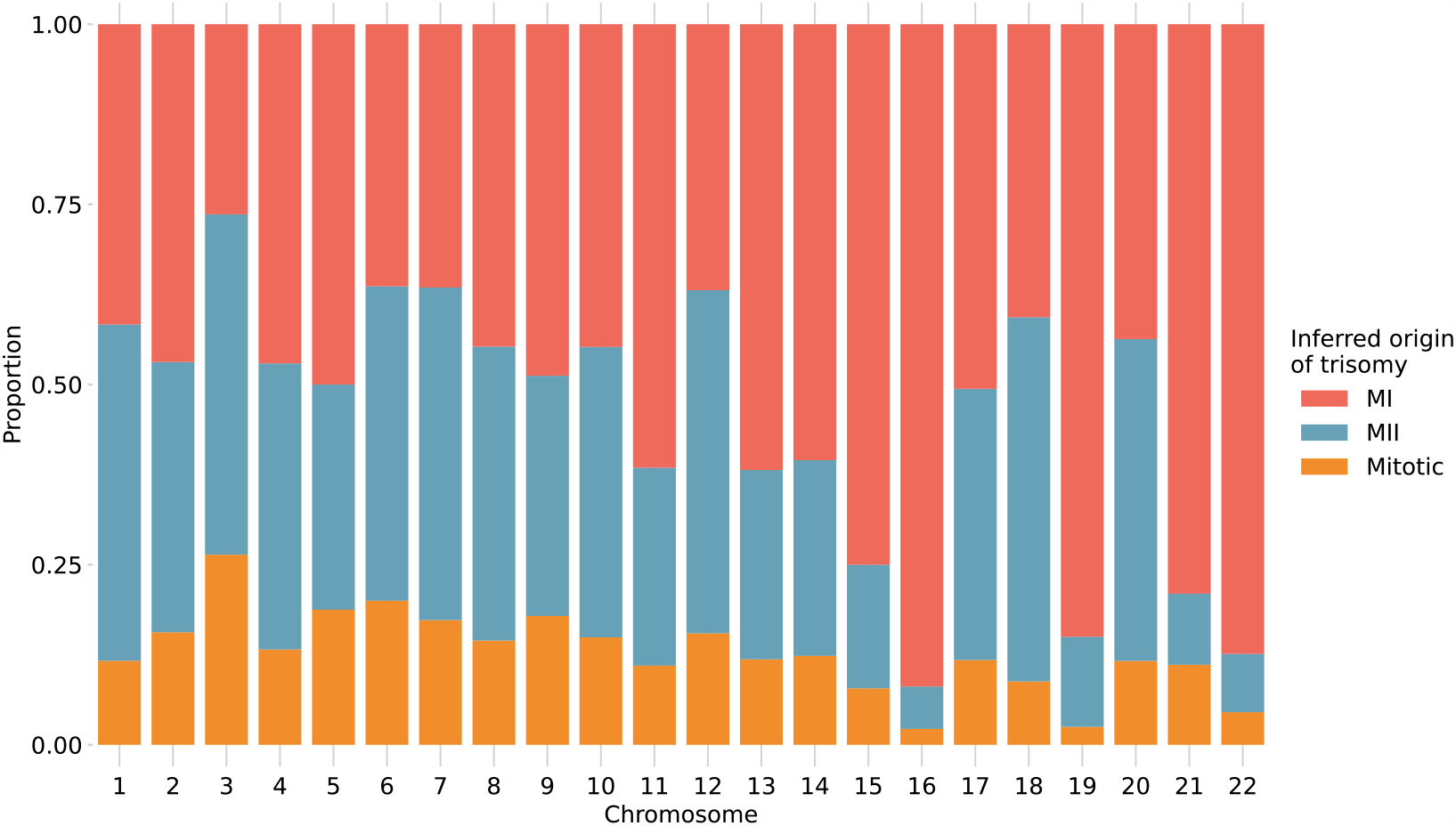
Stratification of trisomies by autosome and inferred source of error. Chromosome-wide patterns of SPH were designated as potentially mitotic in origin (yellow). Samples with tracts of BPH in regions surrounding the centromere were classified as putative MI errors (red), while tracts of BPH elsewhere on the chromosome were classified as putative MII errors (blue).

**Figure 6.**
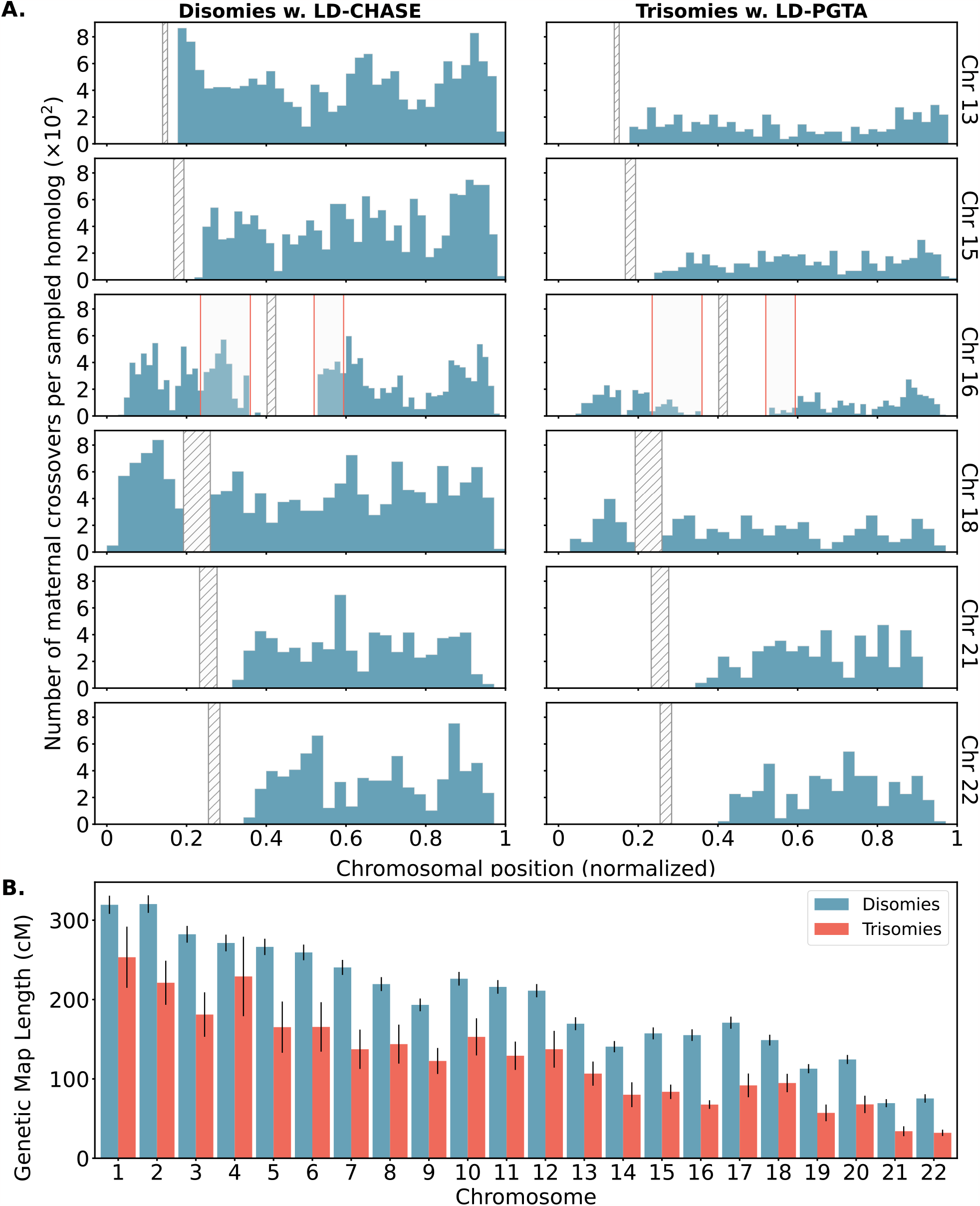
Differences in number and location of inferred crossovers between disomic and trisomic samples. A. Spatial distributions of inferred meiotic crossovers in disomies (left; mapped with LD-CHASE) and trisomies (right; mapped with LD-PGTA) across Chromosomes 13, 15, 16, 18, 21, and 22. Regions with qualitative differences are highlighted in light gray, while centromeres are indicated with diagonal shading. B. Comparisons of female genetic map length for disomies (blue) versus trisomies (red) for each autosome. Observed rates of meiotic crossover were consistently lower for trisomies compared to disomies. Error bars denote 95% confidence intervals as estimated by bootstrapping.

### Ethics approval and consent to participate

The Homewood Institutional Review Board of Johns Hopkins University determined that this work does not qualify as federally-regulated human subjects research (HIRB00011705).

## Data access

Software for implementing our method is available at: https://github.com/mccoy-lab/LD-CHASE. Tables with chromosome abnormalities and crossovers detected for each sample in the study are available in the same repository. Source code and data tables are also uploaded as Supplemental Materials.

## Competing interest statement

D.A. and R.C.M. are co-inventors of the LD-PGTA method, which is the subject of a patent application by Johns Hopkins University.

## Funding

Research reported in this publication was supported by National Institute of General Medical Sciences of the National Institutes of Health under award number R35GM133747. The content is solely the responsibility of the authors and does not necessarily represent the official views of the National Institutes of Health.

## Author contributions

S.M., M.M., S.C., R.A., and C.L. performed data collection and data curation; D.A. and R.C.M. contributed to experimental design and data interpretation; D.A. performed data analysis; D.A. and R.C.M. wrote the paper, with input from S.M. All authors reviewed and approved the manuscript.

## Acknowledgements

Thank you to the staff of Advanced Research Computing at Hopkins.

## Notes

https://github.com/mccoy-lab/LD-CHASE

### Competing Interest Statement

D.A. and R.C.M. are co-inventors of a method used in this study, which is the subject of a patent application by Johns Hopkins University.

### Summary of Updates

Added methods for K-S permutation test.

