## Supplemental Material for "Aberrant landscapes of maternal meiotic crossovers contribute to aneuploidies in human embryos"

### Supplemental Material of the article "Aberrant landscapes of maternal meiotic crossovers contribute to aneuploidies in human embryos" by Ariad et al. (2023)

#### Supplemental Figures

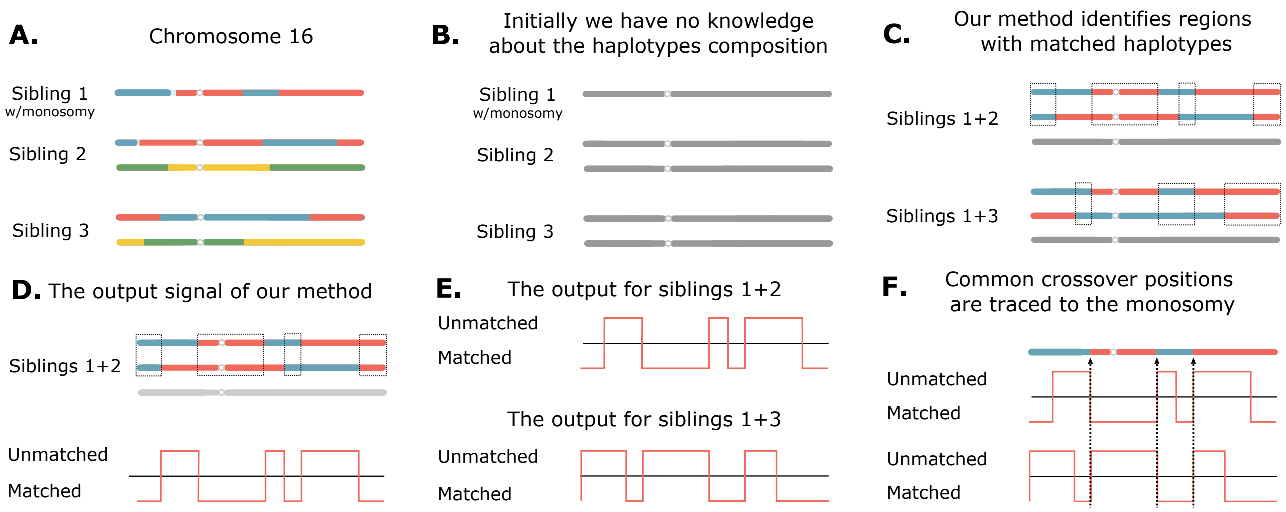


**Figure S1.** *Schematic of our method to detect meiotic crossovers.* **A.** IVF cycles that include a monosomic embryo are identified via conventional coverage-based copy number calling. **B.** Initially, we have no knowledge about the haplotype composition. **C.** For the affected chromosome, each disomic sibling is then contrasted with the monosomic sample to identify regions where the haplotypes match. **D.** Positive signal emitted by our algorithm indicates evidence of non-matching, whereas negative signal indicates evidence of matching. **E.** Output from various sibling embryos is compared in order to attribute crossovers to specific samples. **F.** Apparent crossovers that are shared among sibling embryos can be attributed to the reference monosomic sample, while other crossovers are attributed to the test samples.

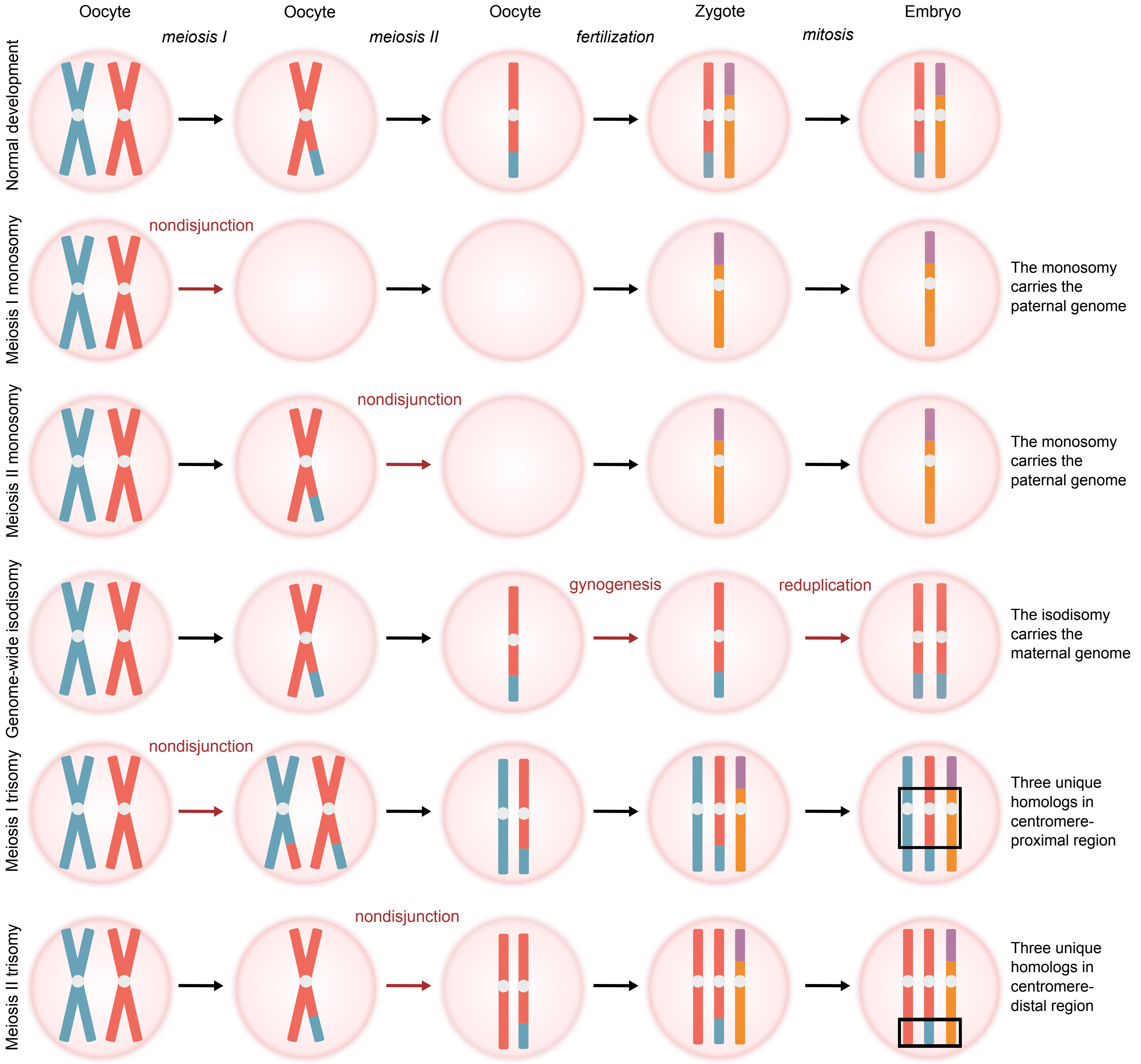


**Figure S2.** *Signatures of various forms of chromosome abnormality with respect to their composition of genetically identical versus distinct parental homologs.* Normal gametogenesis and fertilization produce a zygote with two genetically distinct copies of each chromosome—one copy from each parent. The vast majority of monosomies arise from maternal meiotic errors (occurring either during meiosis I or II), such that the zygote solely possesses the paternally inherited copy of the chromosome. Conversely, in the case of genome-wide uniparental isodisomy (GW-isoUPD), all homologous chromosomes are identical and are typically maternally inherited [33]. Meiotic-origin trisomies may be diagnosed by the presence of one or more tracts with three distinct parental homologs (i.e., transmission of both parental homologs [BPH] from a given parent; indicated by black boxes).

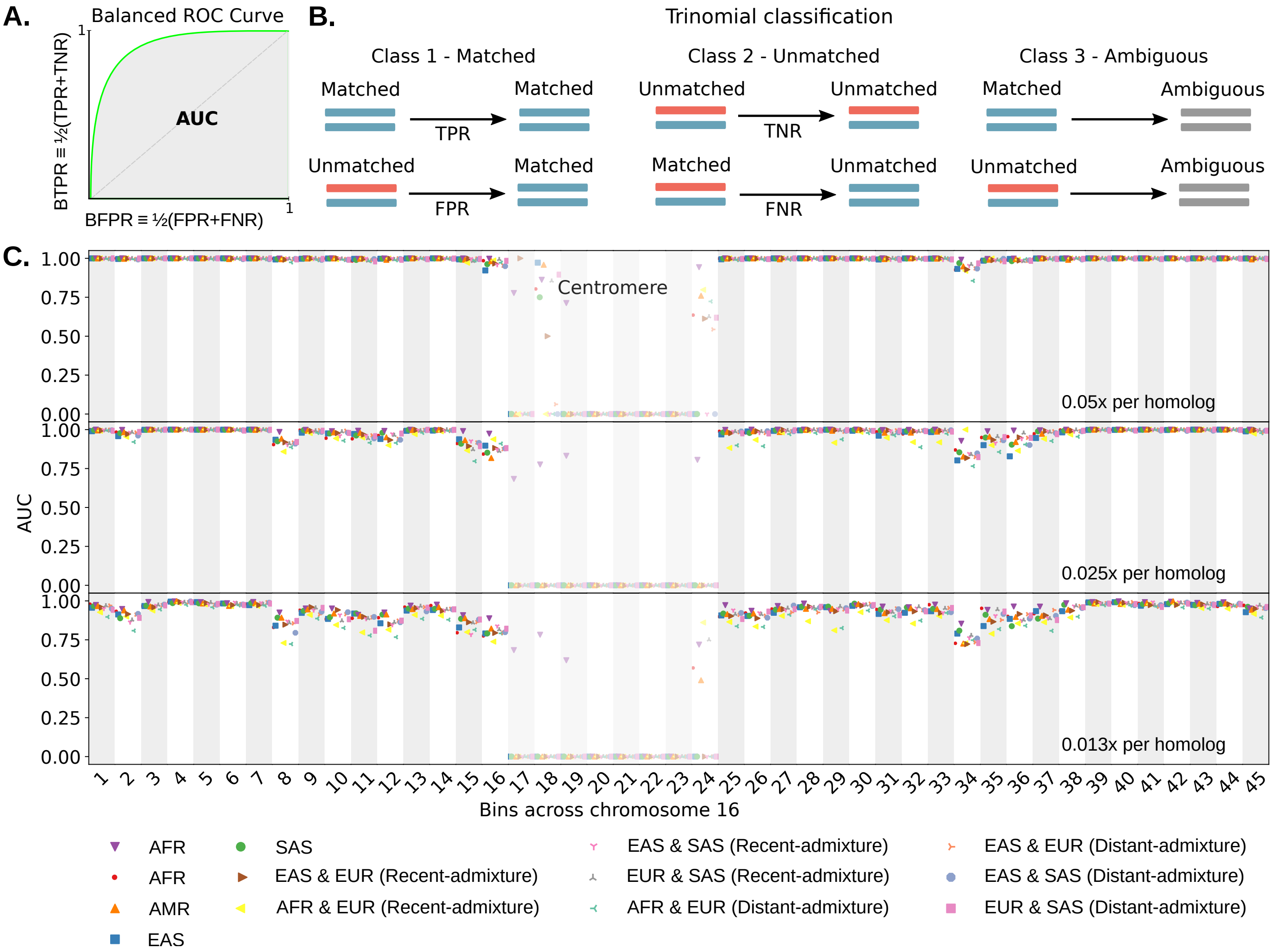


**Figure S3.** *Evaluating the sensitivity and specificity of LD-CHASE along Chromosome 16 based on simulation.* **A.** A generic balanced ROC curve, where the balanced true (false) positive rate is an average of the true (false) positive rate and the true (false) negative rate. **B.** A schematic describing our trinomial classification approach, which includes matched, unmatched, and ambiguous classes. Simulated samples with confidence intervals spanning zero were assigned to the ambiguous class. **C.** Using phased genotypes from the 1000 Genomes Project, we simulated pairs of samples with Monosomy 16 and Disomy 16, where half of the pairs had matched haplotypes and the other half had unmatched haplotypes. Then we divided Chromosome 16 into 45 bins (of $\sim 2$ Mbp). For each bin, we calculated the area under the balanced ROC curve, stratifying over superpopulations from the 1000 Genomes Project and with varying patterns of recent (i.e., first generation) and distant admixture. Balanced ROC curves were calculated by varying the $z$-score threshold.

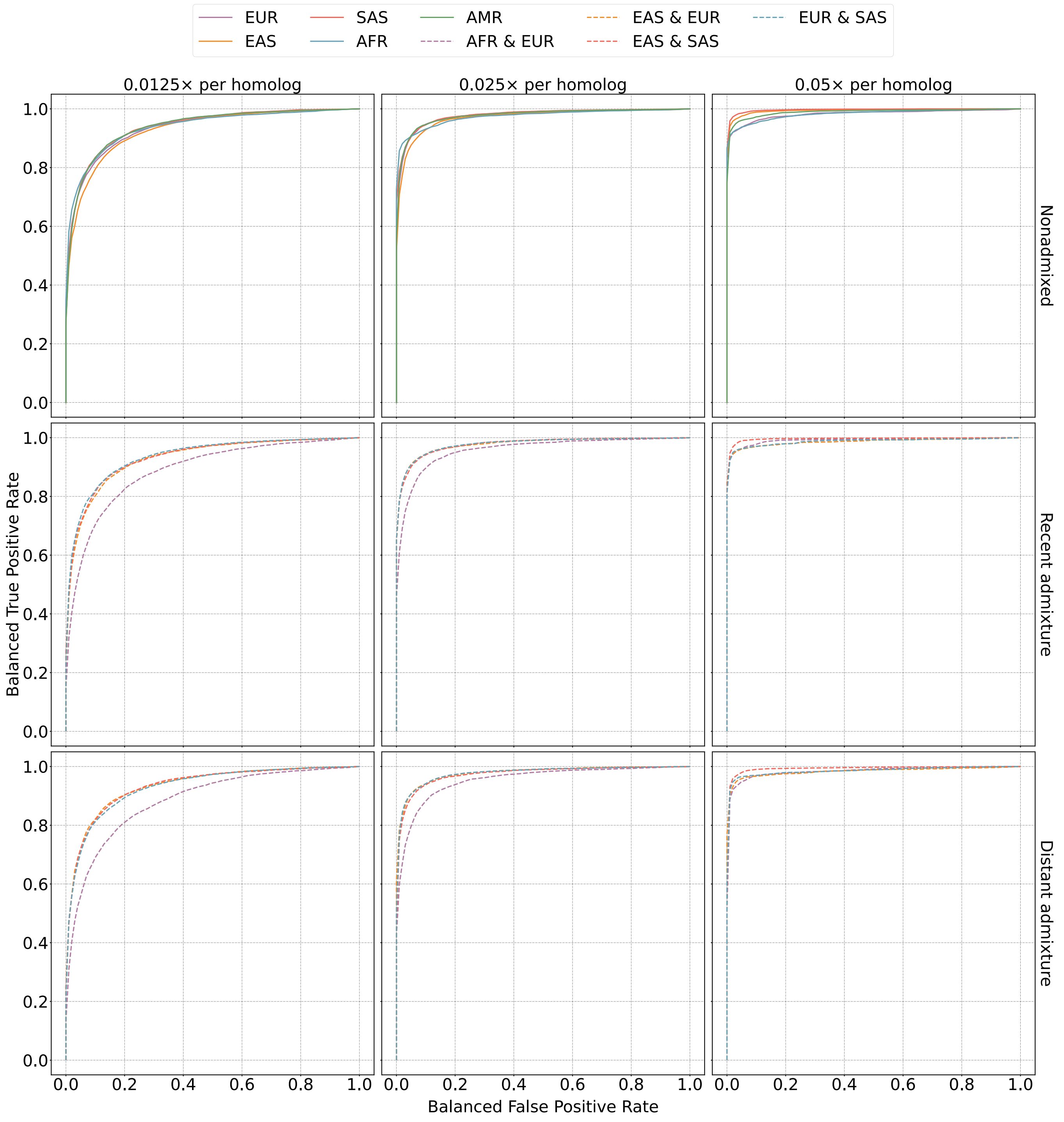


**Figure S4.** *Evaluating the sensitivity and specificity of LD-CHASE across populations based on simulation.* Balanced ROC curves for simulations at varying depths of coverage, sample ancestries, and admixture scenarios, where the balanced true (false) positive rate is an average of the true (false) positive rate and the true (false) negative rate. Using phased genotypes from the 1000 Genomes Project, we simulated pairs of Monosomy 16 and Disomy 16, where half of the pairs had matched haplotypes and the other half had unmatched haplotypes. Then we divided Chromosome 16 into 45 bins (of $\sim$ 2 Mbp). For each bin, we calculated a balanced ROC curve and then averaged the curves across bins by using a linear interpolation to obtain the balanced true positive rate that is associated with a fixed balanced false positive rate. All the simulated admixture involved equal proportions of ancestry from the component populations. Balanced ROC curves were calculated by varying the $z$-score threshold.

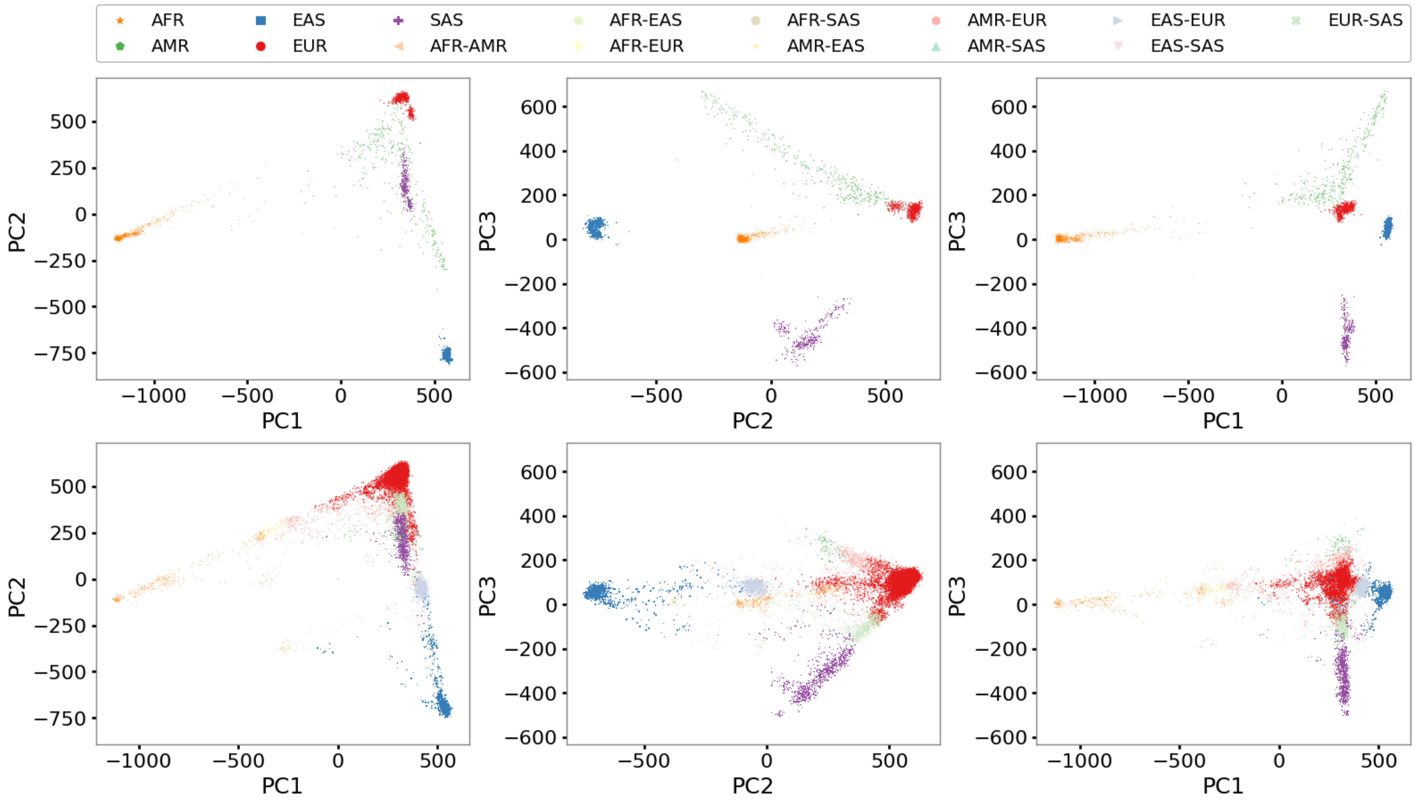


**Figure S5.** *Ancestry inference from low-coverage PGT-A data.* Genetic similarity between test samples and external reference samples informs the selection of ancestry-matched reference panels. Principal component axes were defined based on analysis of 1000 Genomes reference samples and colored according to superpopulation annotations (top row). Low-coverage embryo samples were then projected onto these axes (bottom row) using a Procrustes approach implemented with LASER (v2.0) [59], and genetic similarity to reference samples was determined using the k-nearest neighbors algorithm ($k$ = 150) based on rectilinear distance on the top 32 principal components. For plotting purposes, we associated each test sample with up to two reference superpopulations if a given superpopulation comprised at least 15% of the nearest neighbors.

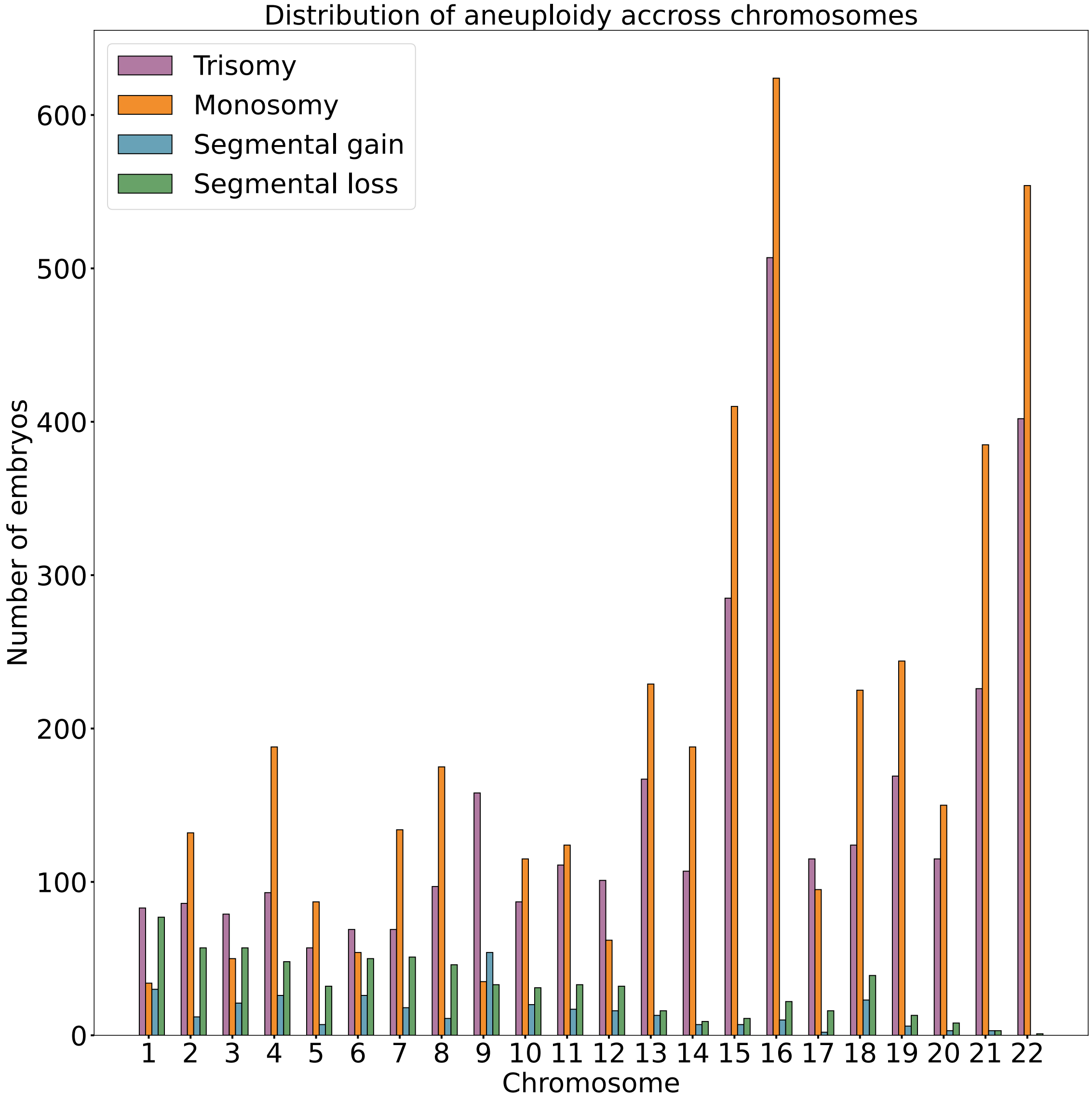


**Figure S6.** *Chromosome-specific counts of aneuploidies detected in PGT-A data.* We analyzed the CReATe PGT-A dataset with WisecondorX to infer the copy number of all autosomes across 20, 114 IVF embryos. We observe that segmental gains and losses are more common on longer chromosomes, while whole-chromosome aneuploidies are more common on short chromosomes, especially Chromosomes 15, 16, 21, and 22, consistent with previous literature.

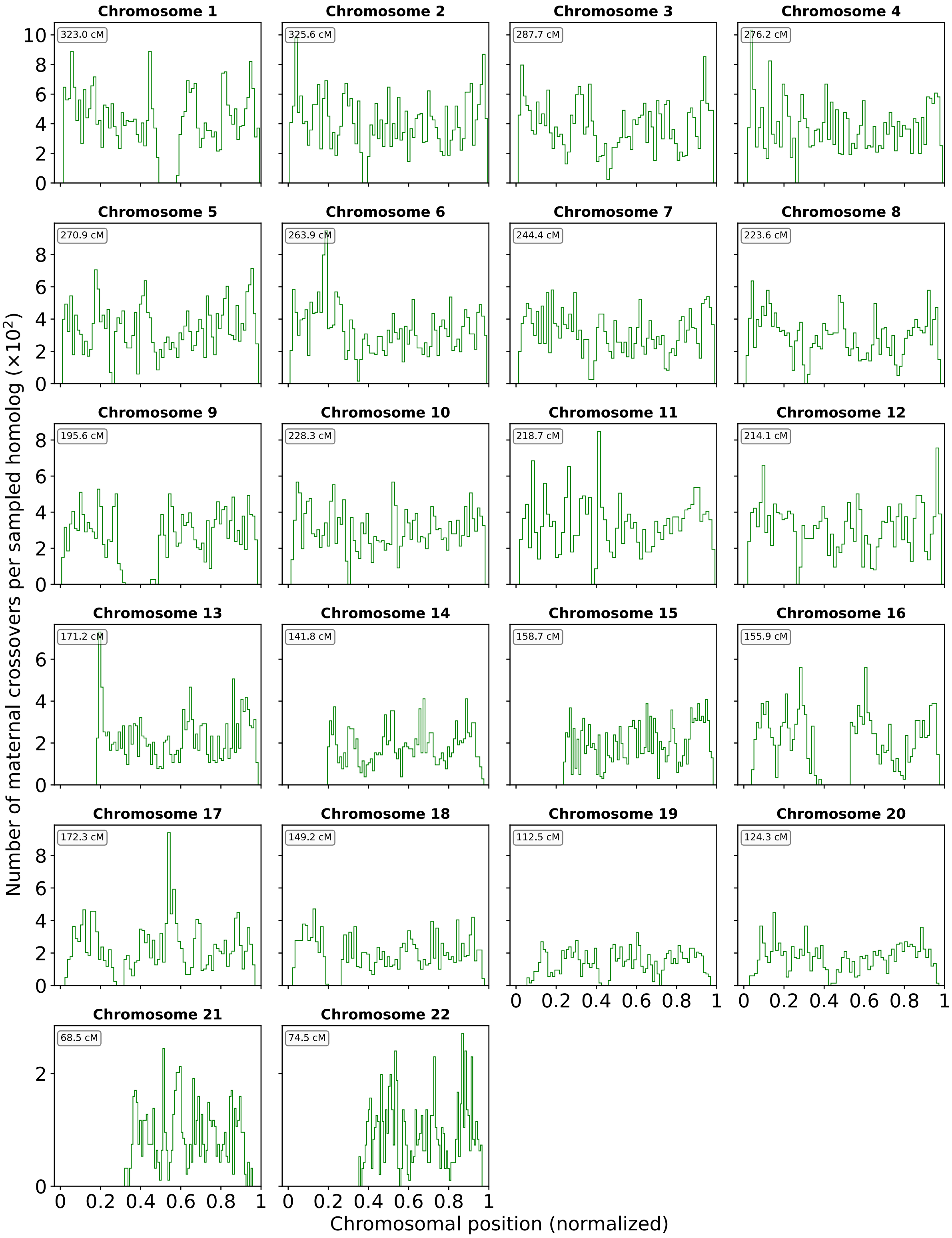


**Figure S7.** *Distribution of maternal crossovers across normal disomic embryo samples.* Crossovers were identified as transitions between tracts that matched versus did not match a sibling haploid/GW-isoUPD sample (only maternal chromosomes present). Each tract was required to cover at least 15 genomic windows and achieve a $z$-score of the at least 1.96. The length of each chromosome is normalized to 1 to aid visualization. The region size, where crossovers occur across sibling embryos at similar chromosomal positions and are solely associated with the reference monosomy, is set to 1.5% of the chromosome size.

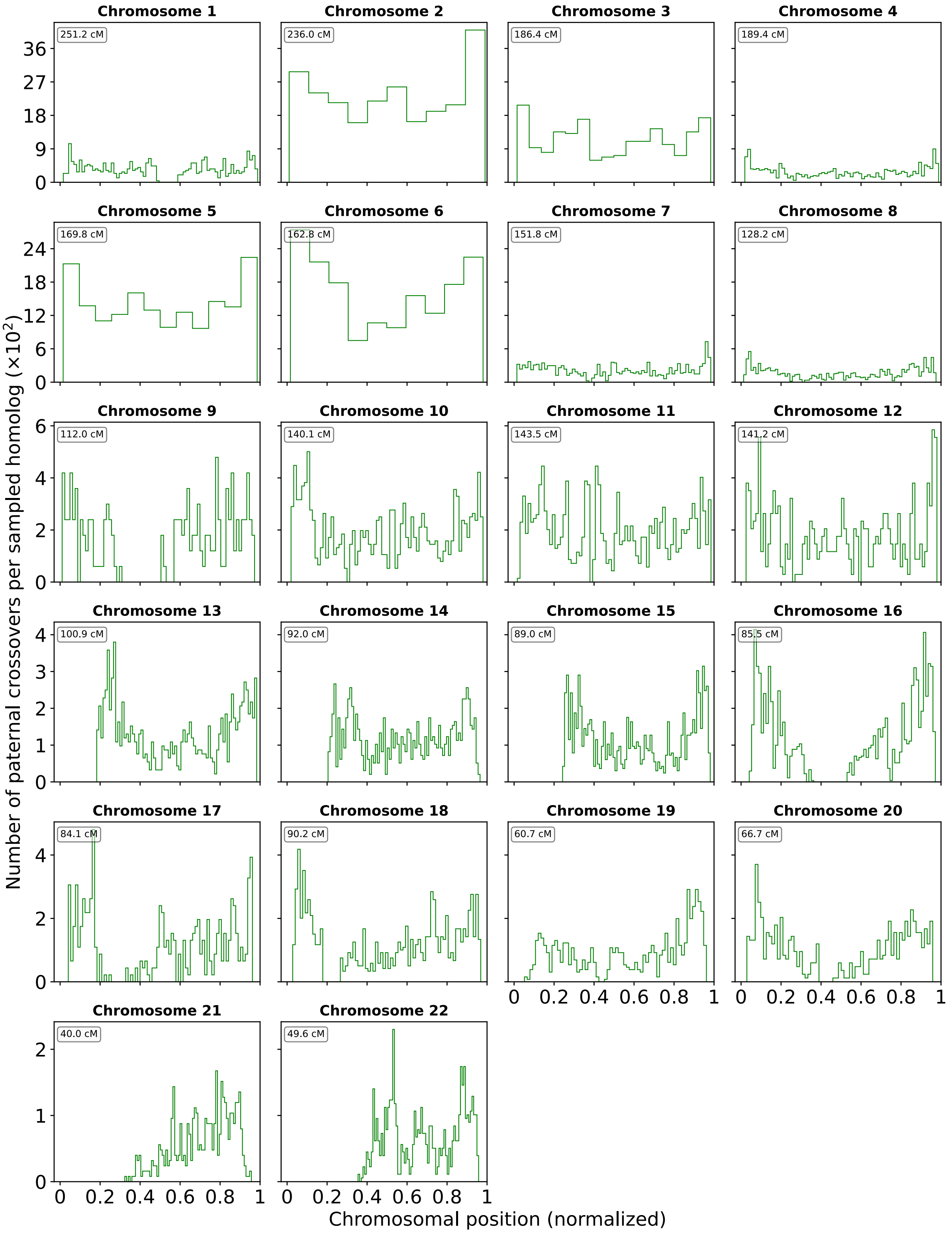


**Figure S8.** *Distribution of paternal crossovers across normal disomic embryo samples.* Crossovers were identified as transitions between tracts that matched versus did not match a sibling monosomic sample (only paternal chromosome present). Each tract was required to cover at least 25 genomic windows and achieve a *z*-score of the at least 1.96. The length of each chromosome is normalized to 1 to aid visualization. The region size, where crossovers occur across sibling embryos at similar chromosomal positions and are solely associated with the reference monosomy, is set to 3% of the chromosome size.

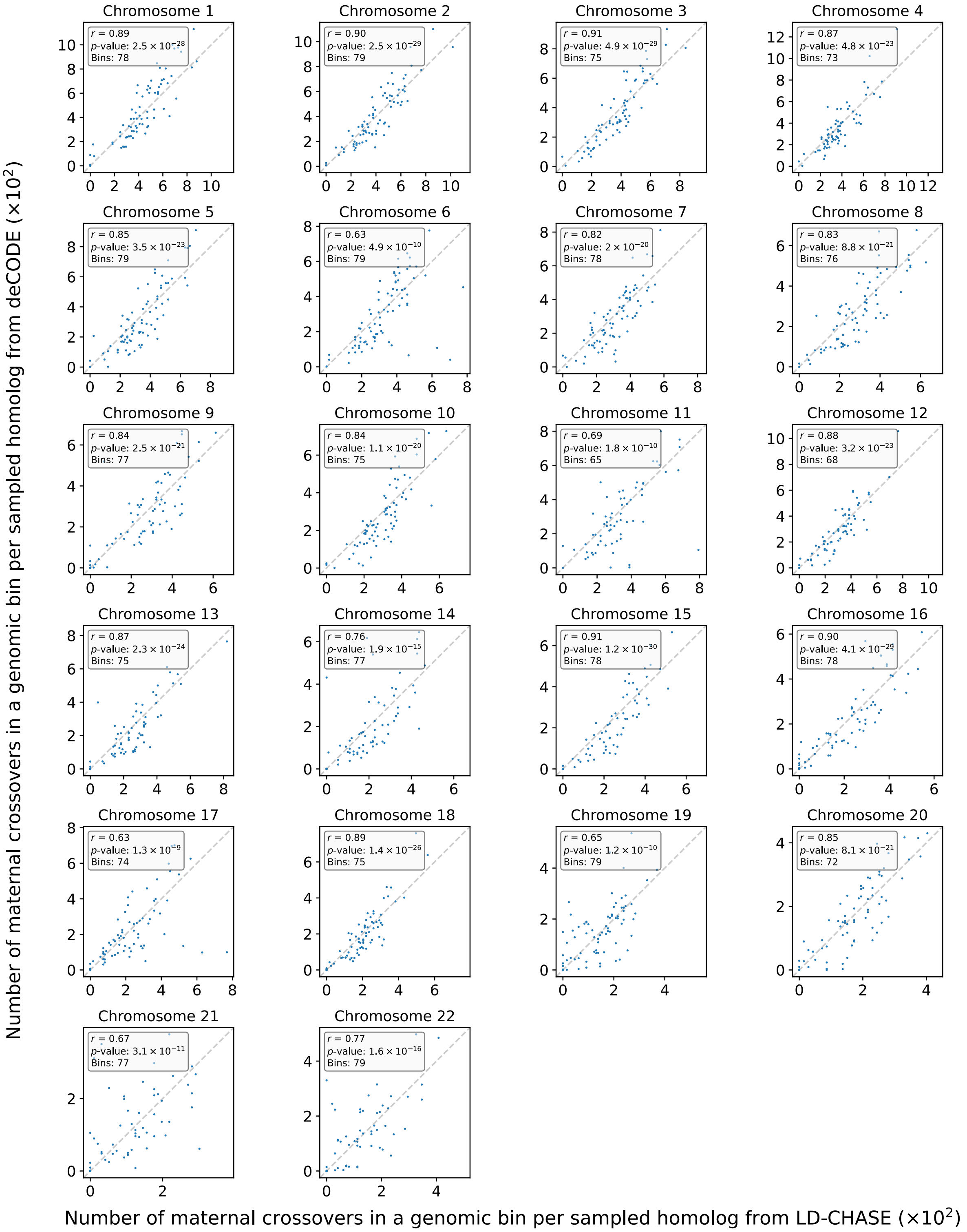


**Figure S9.** *Validating the genomic distribution of maternal crossovers.* We compared rates of maternal crossovers per bin inferred in our study to published data from deCODE [25]. We defined the recombination rate as the number of crossovers in a genomic bin per sampled homolog and computed Pearson correlation coefficients ($r$) between the studies. The number of bins per chromosome was selected to minimize the $p$-value (but not the correlation coefficient). The deCODE recombination map is treated as gold-standard, as maternal crossovers were inferred from 70, 086 samples [25].

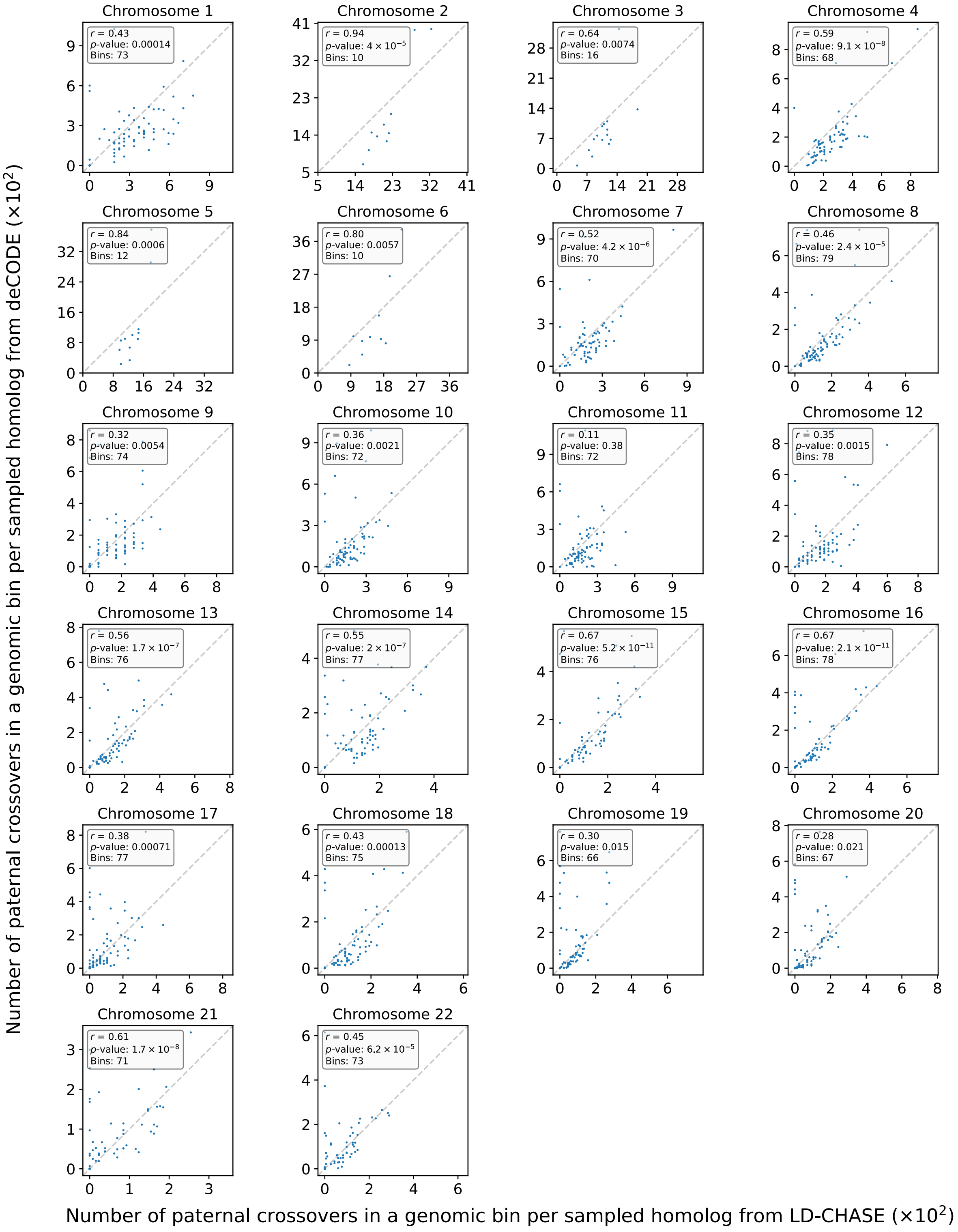


**Figure S10.** *Validating the genomic distribution of paternal crossovers.* We compared rates of paternal crossovers per bin inferred in our study to published data from deCODE [25]. We defined the recombination rate as the number of crossovers in a genomic bin per sampled homolog and computed Pearson correlation coefficients ($r$) between the studies. The number of bins per chromosome was selected to minimize the $p$-value. The deCODE recombination map is treated as gold-standard, as paternal crossovers were inferred from $56,321$ samples [25].

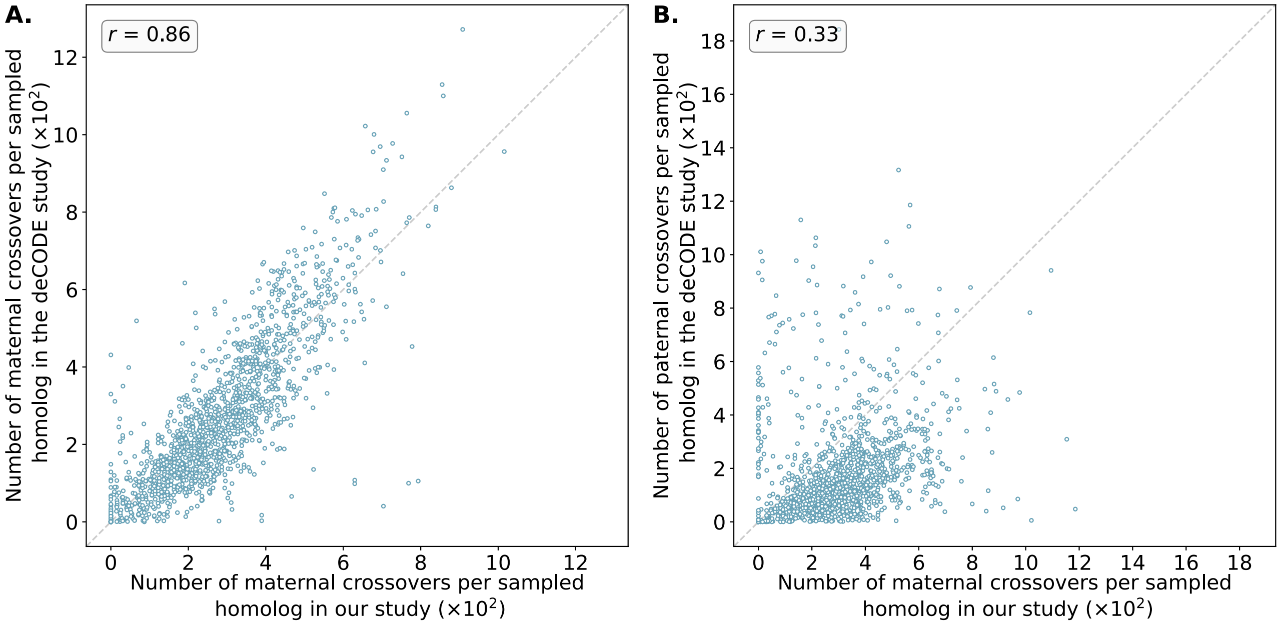


**Figure S11.** *Validating the assumed parental (i.e., sex-specific) origins of crossovers.* **A.** Comparing rates of putative maternal crossovers (identified via comparison to haploid/GW-isoUPD embryos) per bin inferred in our study to published female-specific crossovers from deCODE. **B.** Comparing rates of putative maternal crossovers (identified via comparison to haploid / GW-isoUPD embryos) per bin inferred in our study to published male-specific crossovers from deCODE. The reduction in the correlation coefficient by 0.53 supports our assumption that the vast majority of haploid/GW-isoUPD embryos solely possess the maternally inherited genome, consistent with previous reports [33].

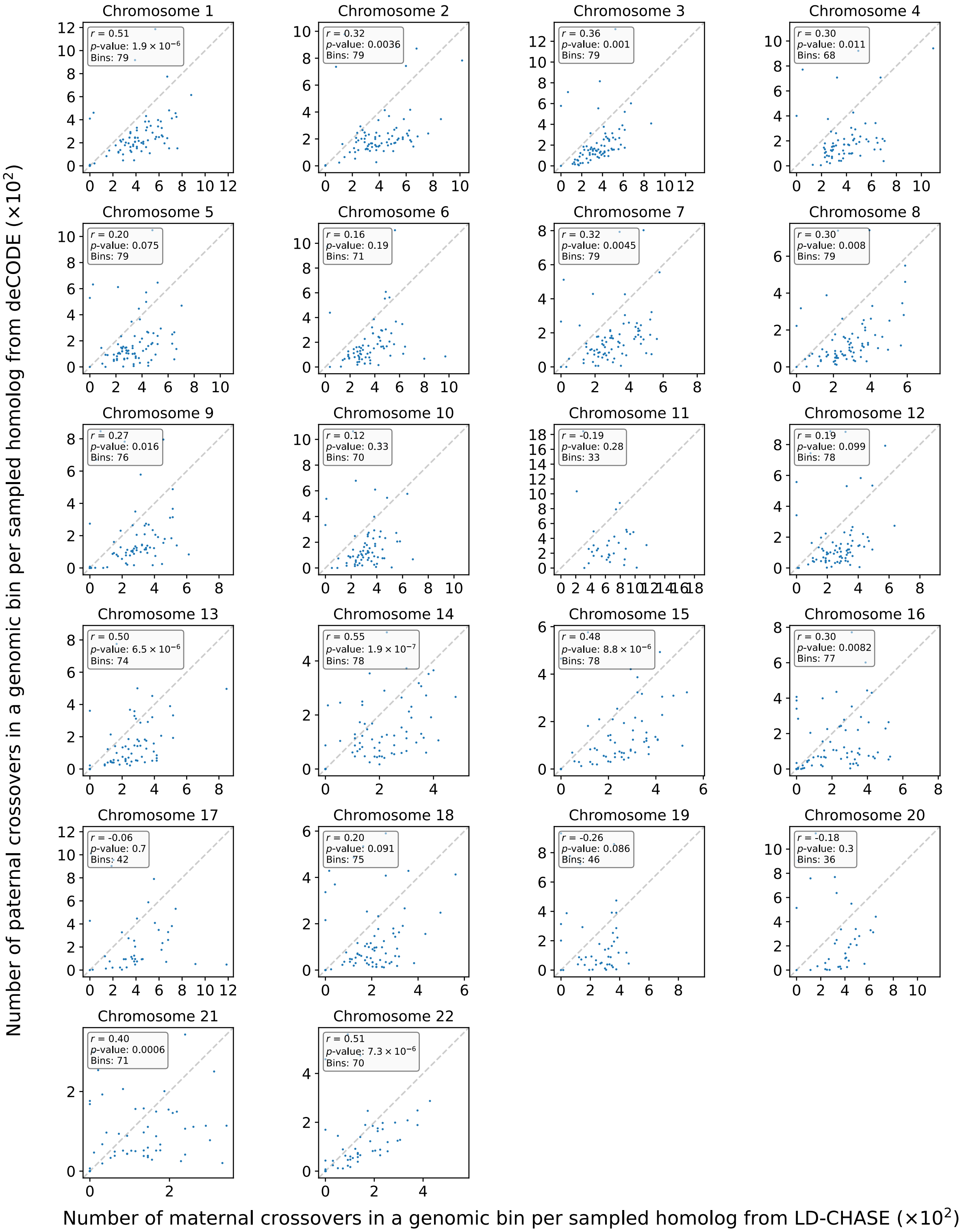


**Figure S12.** *Validating the assumed parental (i.e., sex-specific) origins of crossovers.* We compared rates of putative maternal crossovers (identified via comparison to haploid/GW-isoUPD embryos) per bin inferred in our study to published male-specific crossovers from deCODE, stratifying across autosomal chromosomes. The Pearson correlation ($r$) was reduced by $> 0.25$ for all autosomes, compared to the correlation of our results with deCODE female-specific recombination rate (Figure S9). This supports our hypothesis that vast majority of the effective-haploids carry the maternal genome, consistent with previous reports [33].

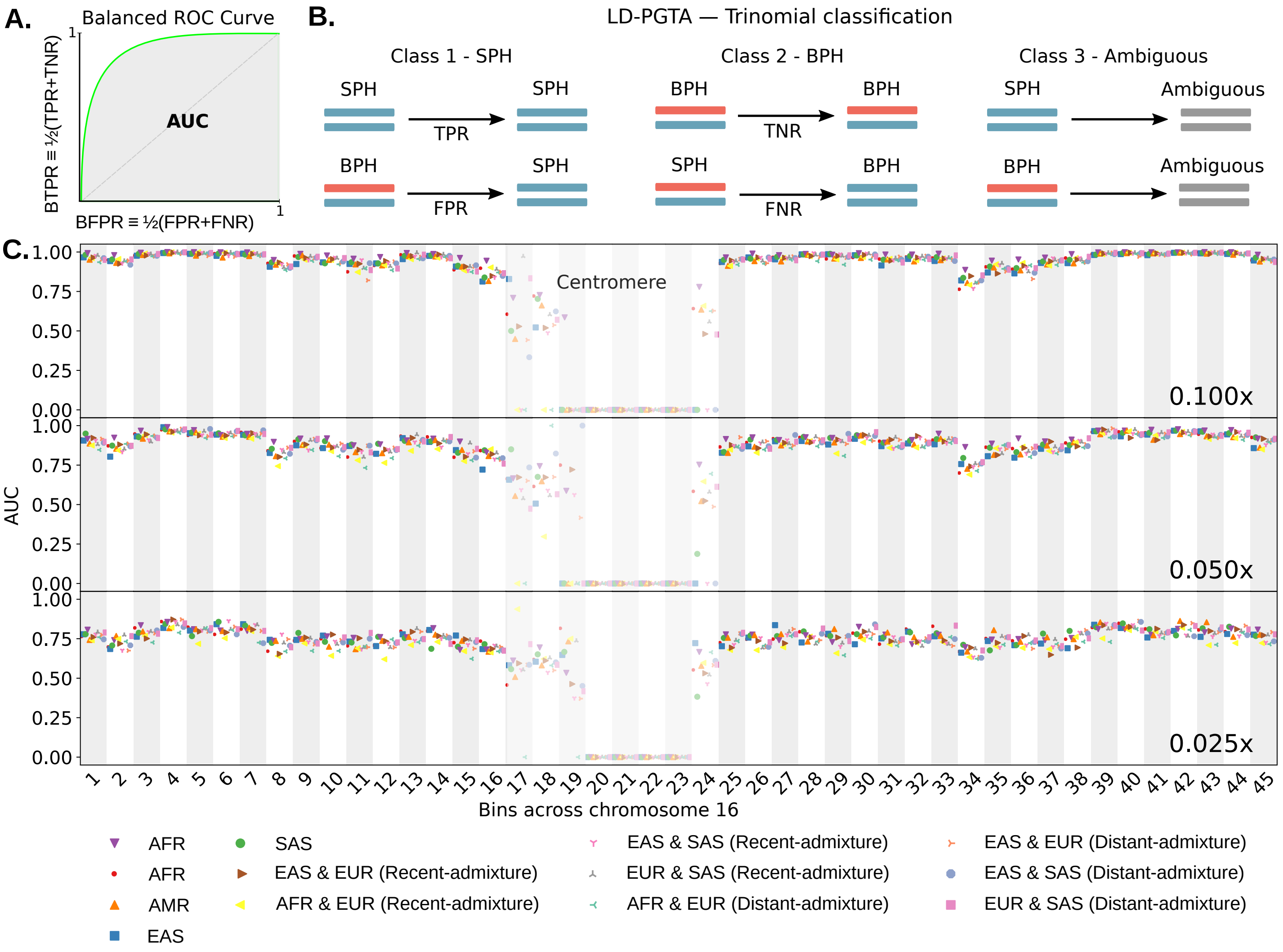


**Figure S13.** *Evaluating the sensitivity and specificity of LD-PGTA [28] along Chromosome 16 based on simulation.* A. A generic balanced ROC curve, where the balanced true (false) positive rate is an average of the true (false) positive rate and the true (false) negative rate. B. A schematic describing our trinomial classification approach, which includes SPH, BPH, and ambiguous classes. Simulated samples with confidence intervals spanning zero were assigned to the ambiguous class. C. Using phased genotypes from the 1000 Genomes Project, we simulated samples with both parental homologs (BPH) and single parental homologs (SPH) configurations for Chromosome 16. Then we divided Chromosome 16 into 45 bins (of $\sim$ 2 Mbp), and for each bin we calculated the area under the balanced ROC curve. Balanced ROC curves were calculated by varying the z-score threshold.

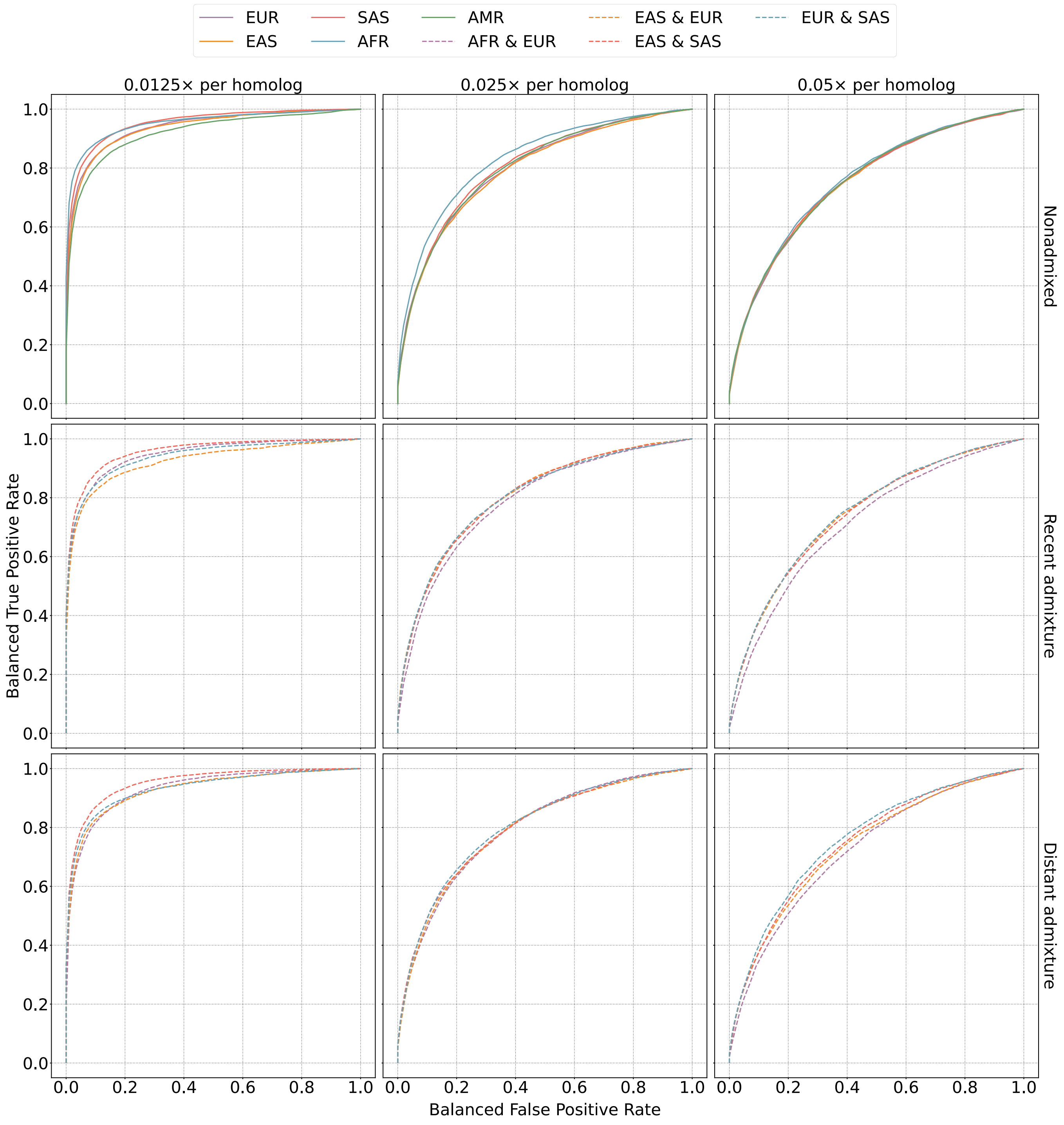


**Figure S14.** *Evaluating the sensitivity and specificity of LD-PGTA [28] across populations based on simulation.* Balanced ROC curves for simulations at varying depths of coverage, sample ancestries, and admixture scenarios, where the balanced true (false) positive rate is an average of the true (false) positive rate and the true (false) negative rate. Using phased genotypes from the 1000 Genomes Project, we simulated samples with both parental homologs (BPH) and single parental homologs (SPH) configurations for Chromosome 16. Then we divided Chromosome 16 into 45 bins (of $\sim 2$ Mbp). For each bin, we calculated a balanced ROC curve and then averaged the curves across bins by using a linear interpolation to obtain the balanced true positive rate that is associated with a fixed balanced false positive rate. All the simulated admixture involved equal proportions of ancestry from the component populations. Balanced ROC curves were calculated by varying the $z$-score threshold.

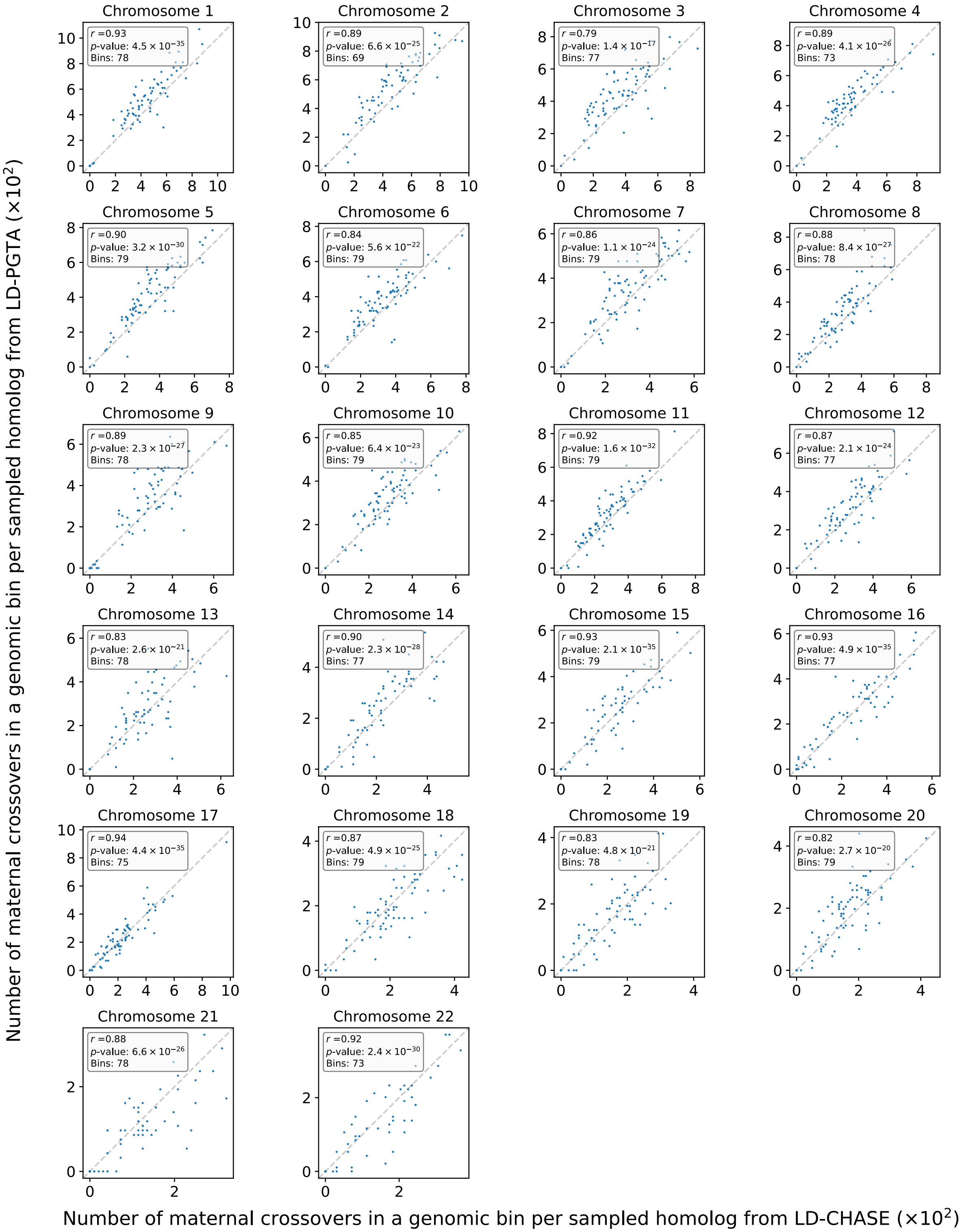


**Figure S15.** *Comparing the distributions of maternal crossovers between LD-CHASE and LD-PGTA.* We calculated the Pearson correlation coefficient between rates of crossovers per genomic bin inferred by LD-PGTA versus LD-CHASE, stratifying across autosomal chromosomes. The number of bins was chosen to minimize the $p$-value (but not the correlation coefficient).

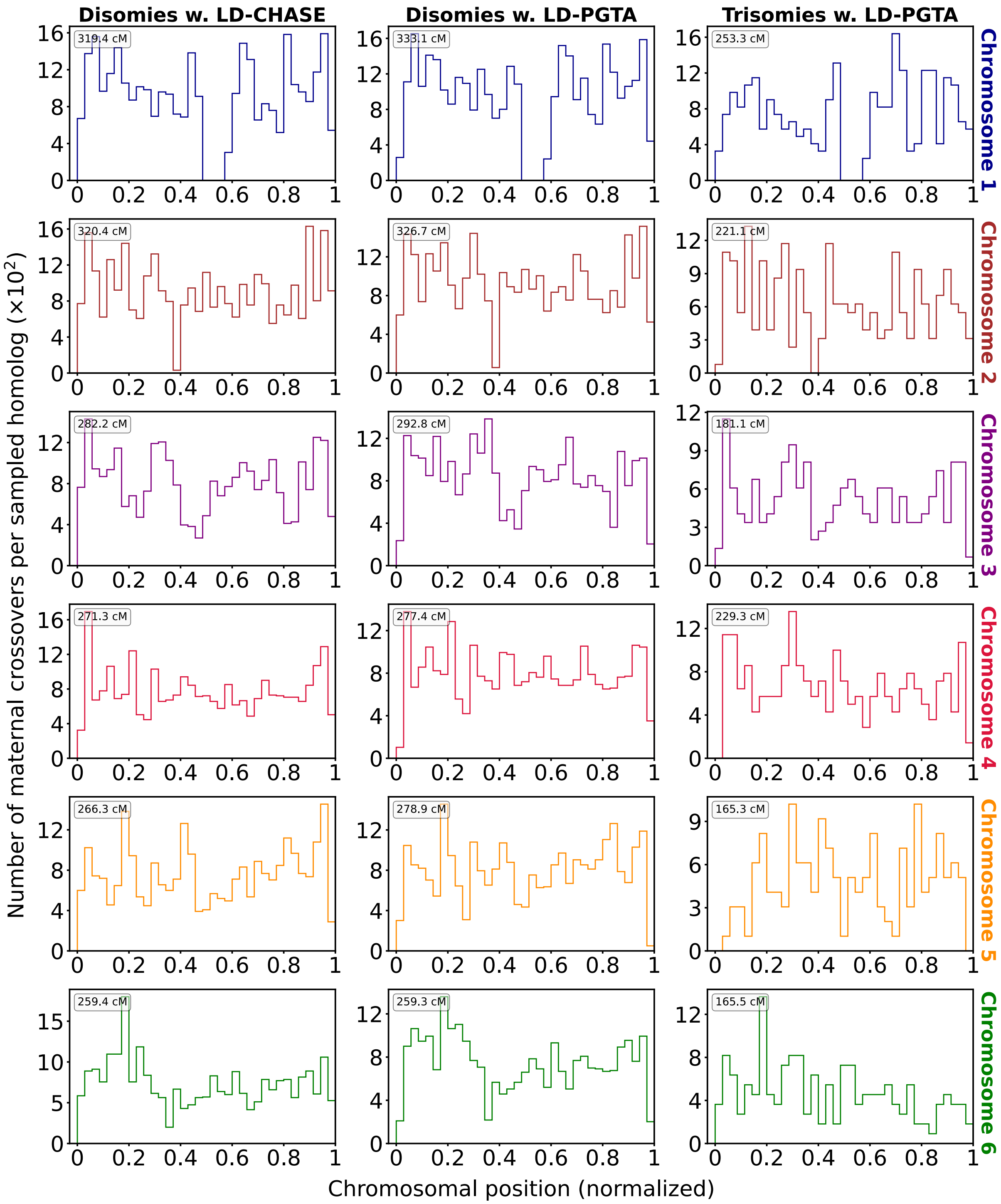


**Figure S16.** *Differences in distributions of crossovers between disomic and trisomic samples across Chromosomes 1 through 6.* We calculated the distribution of maternal crossovers in embryos with two distict homologs (disomy) as well as three distinct homologs (trisomy) for Chromosomes 1-6 (35 bins per chromosome). Crossovers were identified as transitions between regions of "matched" and "unmatched" haplotypes and vice versa, where each region is classified with a $z$-score of the at least 1.96 and included at least 15 genomic windows for LD-CHASE or at least 30 genomic windows for LD-PGTA.

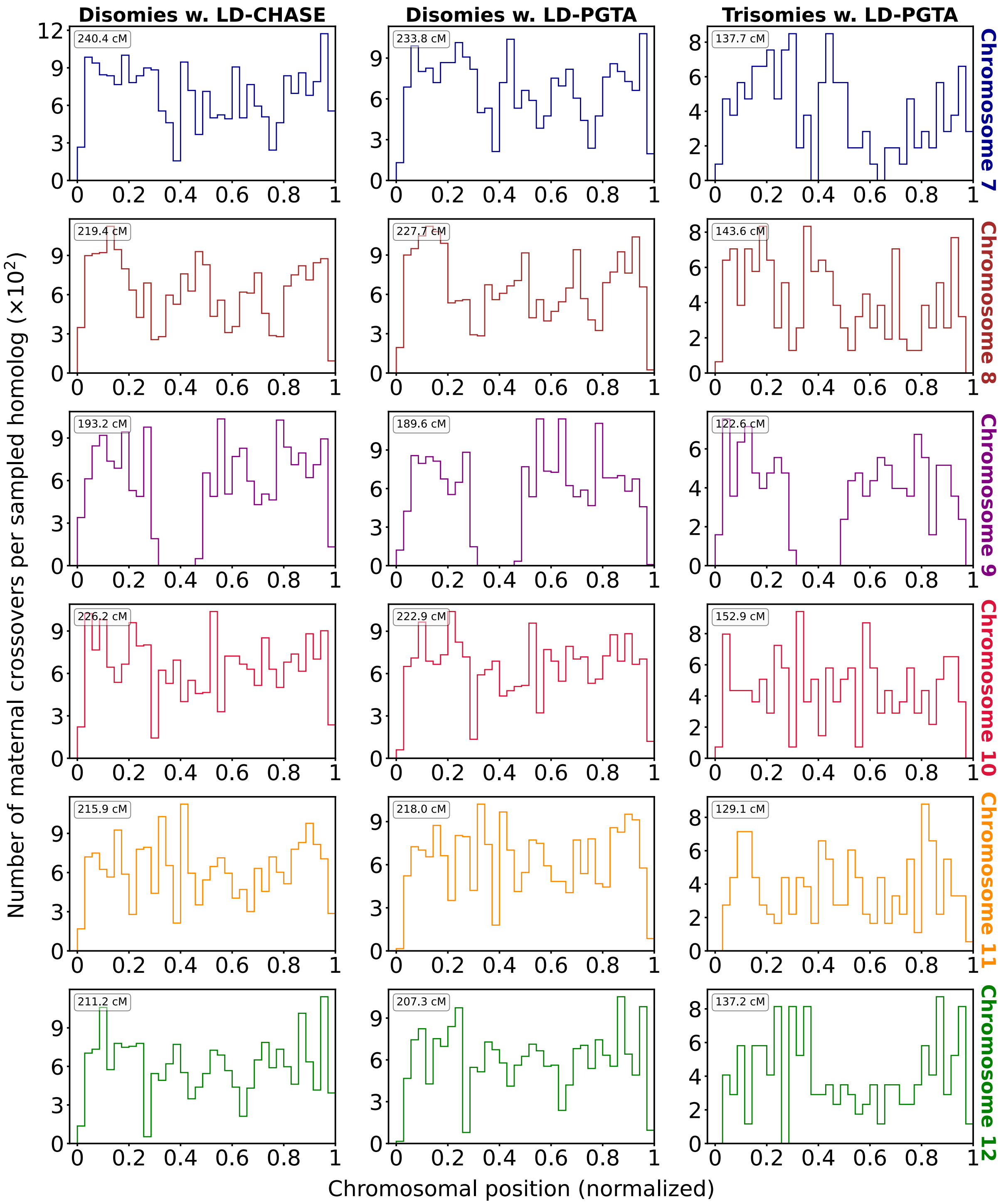


**Figure S17.** *Differences in distributions of crossovers between disomic and trisomic samples across Chromosomes 7 through 12.* We calculated the distribution of maternal crossovers in embryos with two distict homologs (disomy) as well as three distinct homologs (trisomy) for Chromosomes 7-12 (35 bins per chromosome). Crossovers were identified as transitions between regions of "matched" and "unmatched" haplotypes and vice versa, where each region is classified with a $z$-score of the at least 1.96 and included at least 15 genomic windows for LD-CHASE or at least 30 genomic windows for LD-PGTA.

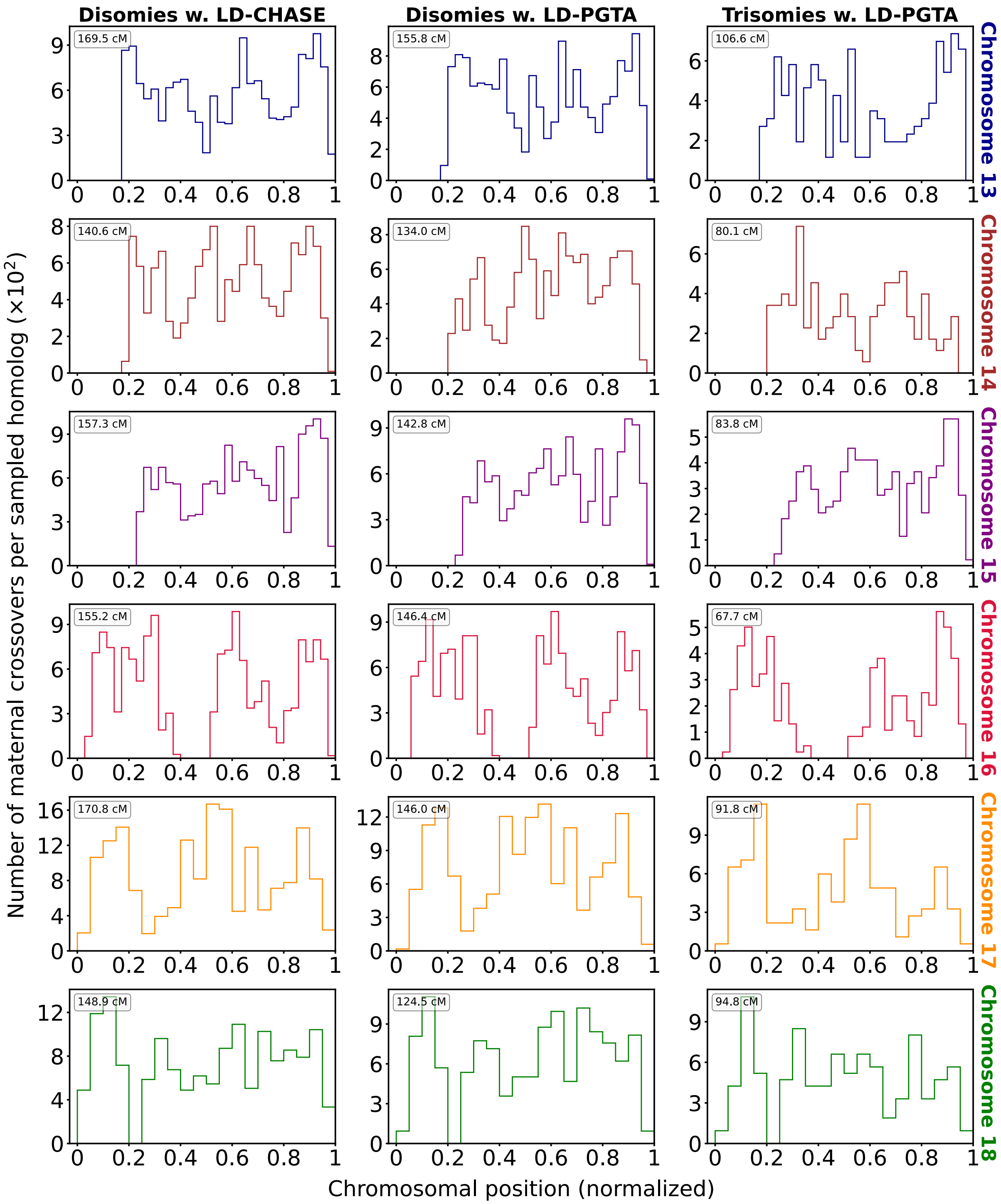


**Figure S18.** *Differences in distributions of crossovers between disomic and trisomic samples across Chromosomes 13 through 18.* We calculated the distribution of maternal crossovers in embryos with two distict homologs (disomy) as well as three distinct homologs (trisomy) for Chromosomes 13-18 (we have 35 bins in Chromosomes 13-16 and 20 bins in Chromosomes 17-18). Crossovers were identified as transitions between regions of "matched" and "unmatched" haplotypes and vice versa, where each region is classified with a $z$-score of the at least 1.96 and included at least 15 genomic windows for LD-CHASE or at least 30 genomic windows for LD-PGTA.

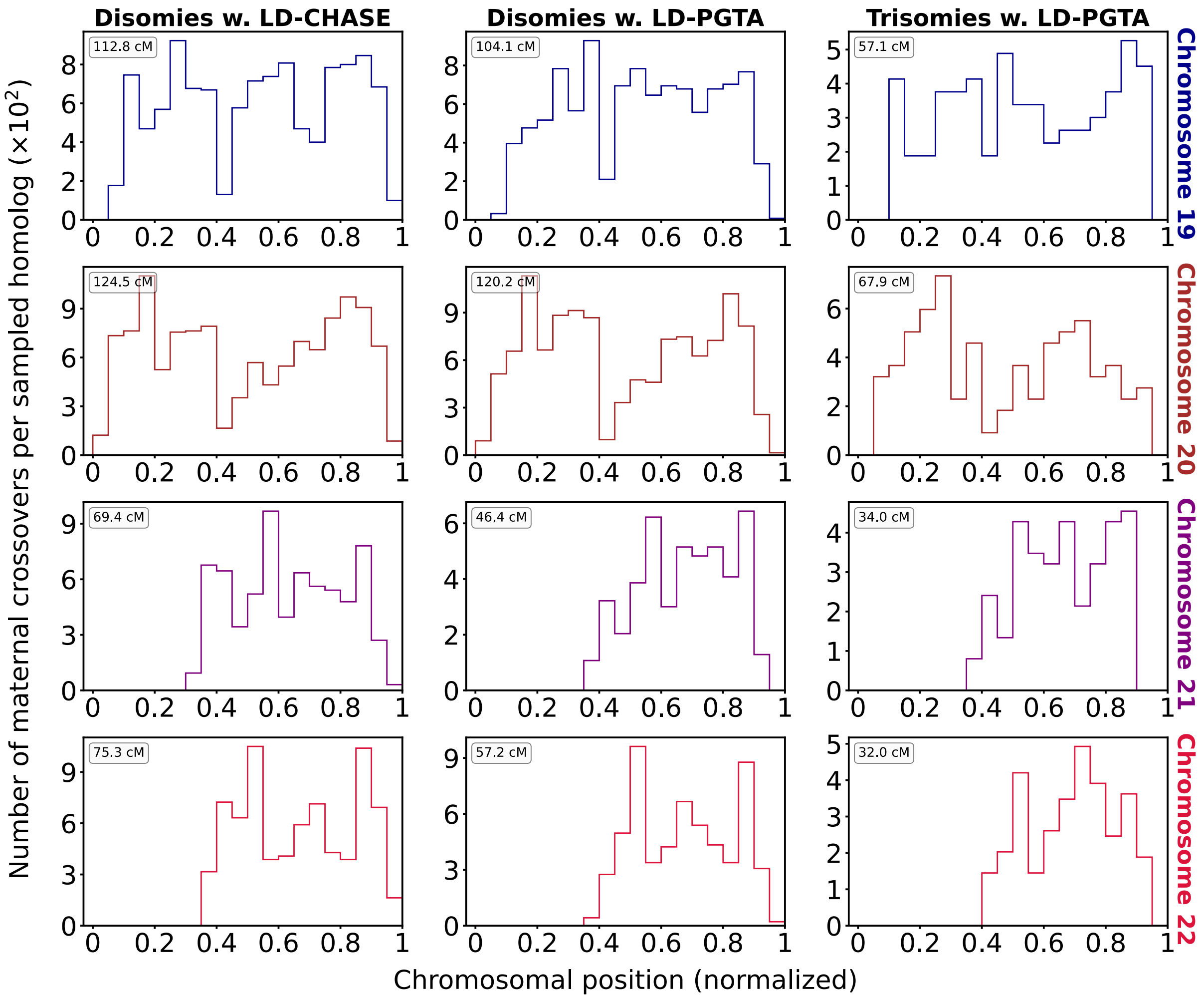


**Figure S19.** *Differences in distributions of crossovers between disomic and trisomic samples across Chromosomes 19 through 22.* We calculated the distribution of maternal crossovers in embryos with two distict homologs (disomy) as well as three distinct homologs (trisomy) for Chromosomes 19-22 (20 bins per chromosome). Crossovers were identified as transitions between regions of “matched” and “unmatched” haplotypes and vice versa, where each region is classified with a $z$-score of the at least 1.96 and included at least 15 genomic windows for LD-CHASE or at least 30 genomic windows for LD-PGTA.

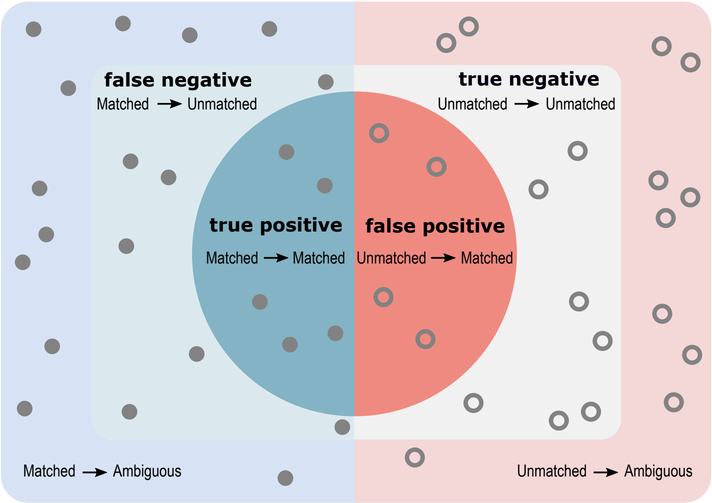


**Figure S20.** A schematic depicting our definitions of true positives, true negatives, false positives, and false negatives with respect to instances of haplotype matching and non-matching among sibling embryo samples. For example, all instances of actual class "unmatched" that were classified as instances of predicted class "matched" are denoted by unmatched⟶ matched. The balanced true (false) positive rate is defined as an average of the true (false) positive and true (false) negative rates.

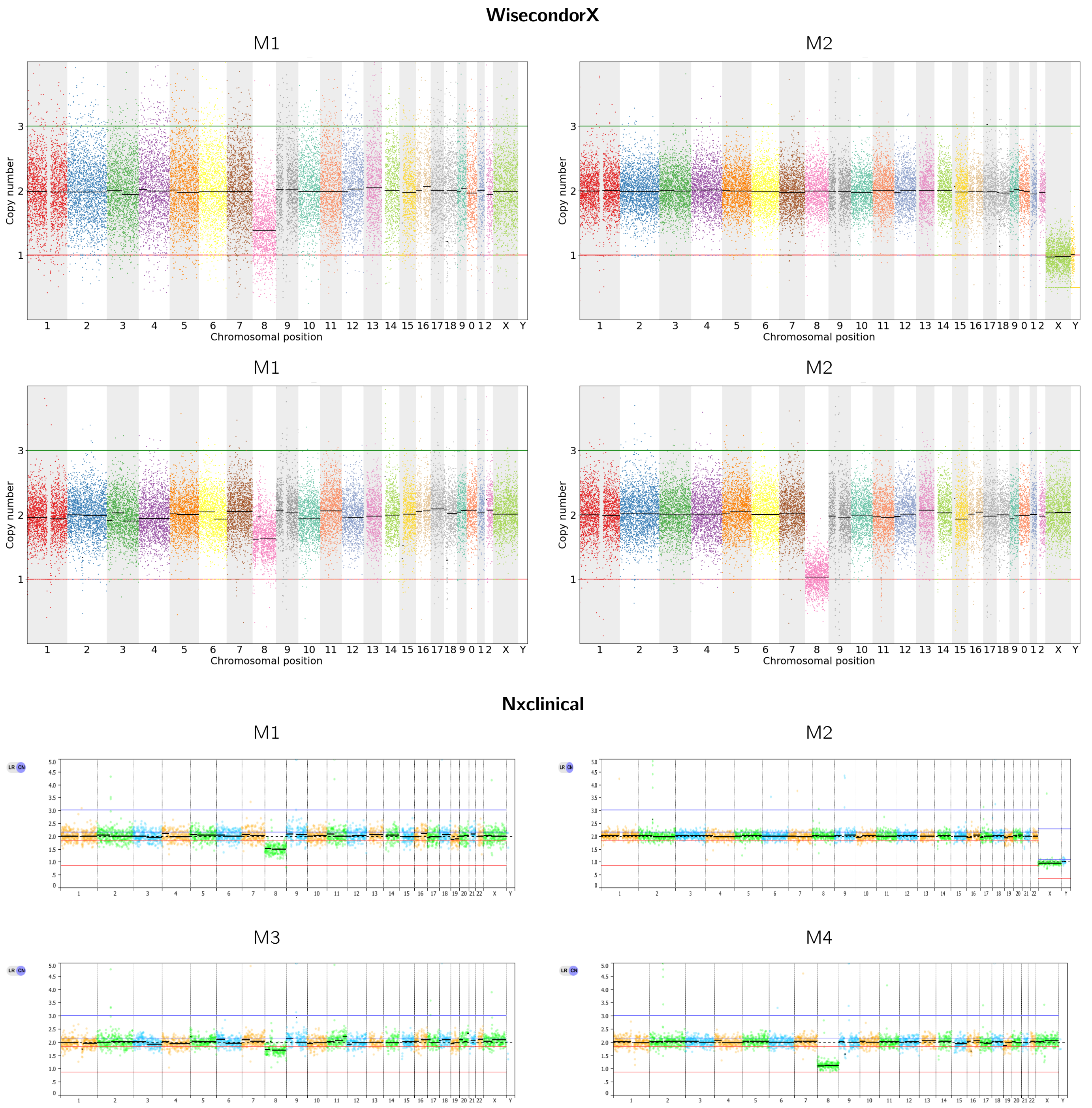


**Figure S21.** *Inferring chromosome copy numbers using conventional coverage-based analysis.* Here we demonstrate that both $N_x$Clinical (a product of Biodiscovery) and WisecondorX produce similar calls for copy number variation of each chromosome. $N_x$Clinical is used for PGT-A by the CReATe Fertility Centre, while WisecondorX was used in this study to analyze the obtained sequences. The bin sizes are 1 Mbp and 100 kbp for $N_x$Clinical and WisecondorX, respectively.

#### Supplemental Tables

| Autosome | Trisomies | Monosomies | Gains | Losses |
|---|---|---|---|---|
| Chromosome 1 | 83 | 34 | 30 | 77 |
| Chromosome 2 | 86 | 132 | 12 | 57 |
| Chromosome 3 | 79 | 50 | 21 | 57 |
| Chromosome 4 | 93 | 188 | 26 | 48 |
| Chromosome 5 | 57 | 87 | 7 | 32 |
| Chromosome 6 | 69 | 54 | 26 | 50 |
| Chromosome 7 | 69 | 134 | 18 | 51 |
| Chromosome 8 | 97 | 175 | 11 | 46 |
| Chromosome 9 | 158 | 35 | 54 | 33 |
| Chromosome 10 | 87 | 115 | 20 | 31 |
| Chromosome 11 | 111 | 124 | 17 | 33 |
| Chromosome 12 | 101 | 62 | 16 | 32 |
| Chromosome 13 | 167 | 229 | 13 | 16 |
| Chromosome 14 | 107 | 188 | 7 | 9 |
| Chromosome 15 | 285 | 410 | 7 | 11 |
| Chromosome 16 | 507 | 624 | 10 | 22 |
| Chromosome 17 | 115 | 95 | 2 | 16 |
| Chromosome 18 | 124 | 225 | 23 | 39 |
| Chromosome 19 | 169 | 244 | 6 | 13 |
| Chromosome 20 | 115 | 150 | 3 | 8 |
| Chromosome 21 | 226 | 385 | 3 | 3 |
| Chromosome 22 | 402 | 554 | 0 | 1 |
| Chromosome $X$ | 21 | 9235 | 0 | 0 |
| Chromosome $Y$ | 0 | 8921 | 0 | 0 |

**Table S1.** *Summarization of chromosome copy number abnormalities that were detected in the CReATe dataset.* Number of whole-chromosome (left two columns) and segmental (right two columns) gains and losses per chromosome as detected by conventional coverage-based analysis.

| Autosome | Meiotic 1 | Meiotic 2 | Mitotic | Ambiguous |
|---|---|---|---|---|
| Chromosome 1 | 25 | 28 | 7 | 1 |
| Chromosome 2 | 30 | 24 | 10 | 0 |
| Chromosome 3 | 19 | 34 | 19 | 2 |
| Chromosome 4 | 32 | 27 | 10 | 1 |
| Chromosome 5 | 24 | 15 | 10 | 0 |
| Chromosome 6 | 20 | 24 | 11 | 0 |
| Chromosome 7 | 19 | 24 | 9 | 1 |
| Chromosome 8 | 34 | 30 | 11 | 3 |
| Chromosome 9 | 60 | 41 | 25 | 0 |
| Chromosome 10 | 30 | 27 | 11 | 1 |
| Chromosome 11 | 56 | 25 | 10 | 0 |
| Chromosome 12 | 31 | 40 | 14 | 1 |
| Chromosome 13 | 73 | 31 | 20 | 5 |
| Chromosome 14 | 49 | 22 | 12 | 5 |
| Chromosome 15 | 153 | 35 | 19 | 12 |
| Chromosome 16 | 375 | 24 | 17 | 3 |
| Chromosome 17 | 43 | 32 | 14 | 3 |
| Chromosome 18 | 37 | 46 | 11 | 12 |
| Chromosome 19 | 102 | 15 | 12 | 4 |
| Chromosome 20 | 45 | 46 | 13 | 5 |
| Chromosome 21 | 128 | 16 | 38 | 5 |
| Chromosome 22 | 270 | 25 | 42 | 8 |

**Table S2.** *Stratification of trisomies by autosome and inferred source of error.* Chromosome-wide patterns of SPH were designated as potentially mitotic in origin.

| Chromosome Number | Disomies w. LD-CHASE vs. Disomies w. LD-PGTA | | Disomies w. LD-CHASE vs. Trisomes w. LD-PGTA | | Disomies w. LD-PGTA vs. Trisomies w. LD-PGTA | |
|---|---|---|---|---|---|---|
| | KS test statistic | *p*-value | KS test statistic | *p*-value | KS test statistic | *p*-value |
| 1 | 0.027 | $1.127 \times 10^{-1}$ | 0.078 | $5.612 \times 10^{-2}$ | 0.069 | $1.299 \times 10^{-1}$ |
| 2 | 0.022 | $2.960 \times 10^{-1}$ | 0.051 | $4.809 \times 10^{-1}$ | 0.052 | $4.583 \times 10^{-1}$ |
| 3 | 0.040 | $6.324 \times 10^{-3}$ | 0.051 | $5.116 \times 10^{-1}$ | 0.035 | $9.070 \times 10^{-1}$ |
| 4 | 0.029 | $1.464 \times 10^{-1}$ | 0.048 | $4.965 \times 10^{-1}$ | 0.056 | $3.059 \times 10^{-1}$ |
| 5 | 0.033 | $5.387 \times 10^{-2}$ | 0.092 | $1.393 \times 10^{-1}$ | 0.086 | $1.898 \times 10^{-1}$ |
| 6 | 0.041 | $8.372 \times 10^{-3}$ | 0.106 | $3.899 \times 10^{-2}$ | 0.113 | $2.242 \times 10^{-2}$ |
| 7 | 0.025 | $3.052 \times 10^{-1}$ | 0.146 | $4.664 \times 10^{-3}$ | 0.140 | $7.652 \times 10^{-3}$ |
| 8 | 0.022 | $4.920 \times 10^{-1}$ | 0.096 | $4.275 \times 10^{-2}$ | 0.102 | $2.504 \times 10^{-2}$ |
| 9 | 0.052 | $5.577 \times 10^{-3}$ | 0.066 | $1.801 \times 10^{-1}$ | 0.072 | $1.141 \times 10^{-1}$ |
| 10 | 0.030 | $1.413 \times 10^{-1}$ | 0.052 | $6.344 \times 10^{-1}$ | 0.059 | $4.885 \times 10^{-1}$ |
| 11 | 0.023 | $4.715 \times 10^{-1}$ | 0.055 | $5.005 \times 10^{-1}$ | 0.049 | $6.573 \times 10^{-1}$ |
| 12 | 0.040 | $2.676 \times 10^{-2}$ | 0.059 | $4.171 \times 10^{-1}$ | 0.064 | $3.149 \times 10^{-1}$ |
| 13 | 0.058 | $7.232 \times 10^{-3}$ | 0.069 | $1.996 \times 10^{-1}$ | 0.076 | $1.240 \times 10^{-1}$ |
| 14 | 0.067 | $3.311 \times 10^{-3}$ | 0.138 | $1.314 \times 10^{-2}$ | 0.181 | $3.745 \times 10^{-4}$ |
| 15 | 0.038 | $2.289 \times 10^{-1}$ | 0.050 | $4.257 \times 10^{-1}$ | 0.033 | $8.897 \times 10^{-1}$ |
| 16 | 0.031 | $4.005 \times 10^{-1}$ | 0.112 | $4.428 \times 10^{-5}$ | 0.104 | $2.270 \times 10^{-4}$ |
| 17 | 0.043 | $6.496 \times 10^{-2}$ | 0.086 | $1.904 \times 10^{-1}$ | 0.086 | $1.957 \times 10^{-1}$ |
| 18 | 0.049 | $4.256 \times 10^{-2}$ | 0.079 | $1.953 \times 10^{-1}$ | 0.115 | $1.751 \times 10^{-2}$ |
| 19 | 0.055 | $3.418 \times 10^{-2}$ | 0.062 | $6.387 \times 10^{-1}$ | 0.086 | $2.480 \times 10^{-1}$ |
| 20 | 0.054 | $1.850 \times 10^{-2}$ | 0.123 | $3.041 \times 10^{-2}$ | 0.071 | $4.783 \times 10^{-1}$ |
| 21 | 0.134 | $1.815 \times 10^{-4}$ | 0.132 | $4.494 \times 10^{-2}$ | 0.069 | $7.164 \times 10^{-1}$ |
| 22 | 0.100 | $4.283 \times 10^{-3}$ | 0.145 | $1.438 \times 10^{-3}$ | 0.107 | $5.147 \times 10^{-2}$ |

**Table S3.** *Comparisons of genomic distributions of crossovers between disomic and trisomic samples.* A Kolmogorov–Smirnov (KS) test was used for quantifying the difference among the three crossover distributions: (1) Disomies that were analyzed via LD-CHASE, (2) Disomies that were analyzed via LD-PGTA, and (3) Trisomies that were analyzed via LD-PGTA. One advantage of the KS test is that it does not require binning, as it quantifies the distance between two empirical cumulative distributions. Moreover, the exact *p*-value was computed using the permutation method. This involved evaluating all possible combinations of assignments of the combined data into two groups of the sizes of the two original samples and computing the KS statistic for each combination. The *p*-value is then the proportion of permutations that result in a KS statistic as extreme as (or more extreme than) the observed statistic. We note that the *p*-values for the comparison of Disomies with LD-CHASE vs. Trisomes with LD-PGTA were equal to or below 0.05 for chromosomes 7, 14, 16, 20, 21, and 22.

| Chromosome | Number of cycles with at least one monosomy and disomy | Average number of disomic siblings per cycle |
|---|---|---|
| 1 | 33 | 10.15 |
| 2 | 126 | 8.29 |
| 3 | 47 | 7.94 |
| 4 | 170 | 8.68 |
| 5 | 80 | 9.4 |
| 6 | 53 | 9.15 |
| 7 | 125 | 9.24 |
| 8 | 165 | 8.85 |
| 9 | 31 | 10.29 |
| 10 | 103 | 10.0 |
| 11 | 112 | 8.09 |
| 12 | 60 | 7.35 |
| 13 | 214 | 8.33 |
| 14 | 171 | 9.16 |
| 15 | 351 | 7.64 |
| 16 | 529 | 7.41 |
| 17 | 86 | 8.8 |
| 18 | 198 | 8.89 |
| 19 | 215 | 7.38 |
| 20 | 136 | 7.74 |
| 21 | 332 | 7.33 |
| 22 | 470 | 7.08 |

**Table S4.** *Number of IVF cycles used for mapping paternal crossovers.* Relevant cycles are defined as those with at least one embryo affected by monosomy (of one or few chromosomes), as well as at least one embryo that is disomic for the same chromosome. In addition, we report the average number of disomic sibling embryos within these cycles.

| Chromosome | Number of cycles with at least one haploid and disomy | Average number of disomic siblings per cycle |
|---|---|---|
| 1 | 198 | 9.29 |
| 2 | 196 | 9.30 |
| 3 | 197 | 9.38 |
| 4 | 198 | 9.22 |
| 5 | 198 | 9.24 |
| 6 | 201 | 9.19 |
| 7 | 197 | 9.19 |
| 8 | 203 | 9.00 |
| 9 | 199 | 9.26 |
| 10 | 195 | 9.44 |
| 11 | 199 | 9.29 |
| 12 | 198 | 9.35 |
| 13 | 207 | 8.77 |
| 14 | 202 | 9.01 |
| 15 | 199 | 8.95 |
| 16 | 203 | 8.65 |
| 17 | 203 | 9.01 |
| 18 | 203 | 8.99 |
| 19 | 205 | 8.64 |
| 20 | 202 | 9.06 |
| 21 | 205 | 8.72 |
| 22 | 204 | 8.65 |

**Table S5.** *Number of IVF cycles used for mapping maternal crossovers.* Relevant cycles are defined as those with at least one embryo affected by haploidy/GW-isoUPD, as well as at least one embryo that is disomic for one or more chromosomes. In addition, we report the average number of disomic sibling embryos within these cycles.

### Supplemental Methods

#### The “diploid” statistical model for admixture in the previous generation

In this section we derive the a statistical model for reads that are sampled for genomic library of a diploid. Each read that was sampled from the genomic library might have originated from one of two chromosomal copies, denoted as “homolog $f$” and “homolog $g$”. We first consider a particular configuration of 6 reads in which reads A,B,D and F are associated with homolog $f$ and reads C and E are associated with homolog $g$, as shown in Figure S22.

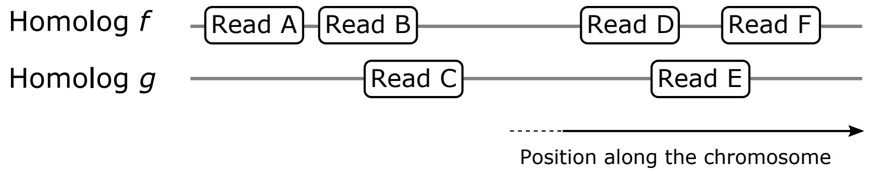


**Figure S22. A possible configuration of 6 reads.** In this configuration, reads A,B,D and F are associated with homolog $f$ and reads C and E are associated with homolog $g$.

The combination of reads that is associated with homologs $f$ and $g$ can be expressed as $\{\mathbf{A}, \mathbf{B}, \mathbf{D}, \mathbf{F}\}$ and $\{\mathbf{C}, \mathbf{E}\}$, respectively. Each read is presented as a sequence of observed alleles at known SNP positions that overlap with the read. Moreover, each allele is presented as a sequence of position and nucleotide, e.g., (821,A). The reads in our example configuration are expressed as:

$$\begin{array}{ll} \mathbf{A}=\big((530,\text{A}),(537,\text{T})\big) & \mathbf{D}=\big((821,\text{A}),\big) \\ \mathbf{B}=\big((641,\text{G}),\big) & \mathbf{E}=\big((957,\text{A}),\big) \\ \mathbf{C}=\big((734,\text{T}),\big) & \mathbf{F}=\big((1001,\text{C}),(1039,\text{T})\big) \end{array}$$

Another useful way to present the reads that are associated with a certain homolog is a sequence of “occupation numbers”: $(n_\text{A}, n_\text{B}, \ldots, n_i, \ldots n_\text{F})$. Here $n_i = 1$ when read $i$ (with $i =$ A,B,. . . ,F) originated from the considered homolog and $n_i = 0$ when the read originated from another homolog. Continuing with the considered configuration, the “occupation numbers” for homologs $f$ and $g$ are shown in Table S6.

| | Read A | Read B | Read C | Read D | Read E | Read F |
|---|---|---|---|---|---|---|
| Homolog $f$ | 1 | 1 | 0 | 1 | 0 | 1 |
| Homolog $g$ | 0 | 0 | 1 | 0 | 1 | 0 |

**Table S6.** *An “occupation table” for a configuration of 6-reads.* Fields with ones mean that the reads are associated with the homologs, while fields with zeros mean the opposite. Thus, the “occupation number representation” of homologs $f$ and $g$ is $(1, 1, 0, 1, 0, 1)$ and $(0, 0, 1, 0, 1, 0)$, respectively.

Next, we would like to calculate the probability of observing such haplotypes as reflected by the 6-reads configuration: $f(\mathbf{A}, \mathbf{B}, \mathbf{D}, \mathbf{F}) \times g(\mathbf{C}, \mathbf{E})$, where $f(\mathbf{A}, \mathbf{B}, \mathbf{D}, \mathbf{F}) \equiv f(1, 1, 0, 1, 0, 1)$ and $g(\mathbf{C}, \mathbf{E}) \equiv g(0, 0, 1, 0, 1, 0)$ are the joint frequencies of the haplotypes $\big((530, A), (537, T), (641, G), (821, A), (1001, C), (1039, T)\big)$ and $\big((734, T), (957, A)\big)$, respectively. Since each homolog originated from a different ancestral population, two population-specific reference panels are required. Using the reference panels in Tables S7 and S8 we deduce that $f(\mathbf{A}, \mathbf{B}, \mathbf{D}, \mathbf{F}) = 1/6$ and $g(\mathbf{C}, \mathbf{E}) = 1/3$.

Based on this knowledge, we are in a position to derive the statistical model. There are $2^6$ possible ways to distribute 6 reads between two homologs, and we assume that all possible configurations of reads are equally likely when sampling from a genomic library. Given a particular configuration of reads, the probability of observing the two haplotypes that are formed by these combinations of reads is $f(\text{Reads} \in \text{Homolog } f) \times g(\text{Reads} \in \text{Homolog } g)$. Thus, the probability of sampling $n$-reads from genomic library of a diploid organism is

$$P(\mathbf{A} \wedge \mathbf{B} \ldots) = \frac{1}{2^n} \sum_{x \in (\mathbb{Z}_2)^n} f(x) g\big(x \text{ XOR } (1, 1, \ldots, 1)\big), \tag{20}$$

| SNP position | Genotypes of individual 1 | | Genotypes of individual 2 | | Genotypes of individual 3 | |
|---|---|---|---|---|---|---|
| 530 | A | C | **A** | A | C | A |
| 537 | C | T | **T** | C | T | T |
| 651 | A | A | **G** | G | G | A |
| 821 | A | A | **A** | G | G | A |
| 1001 | C | T | **C** | C | C | T |
| 1039 | T | A | **T** | A | A | T |

**Table S7.** A population-specific reference panel of individual genotypes, sharing the same ancestral population as homolog $f$. The reference panel describes the genotypes of 3 individual at SNP positions, with the haplotype that was observed in reads **A**,**B**,**D** and **F** being highlighted.

| SNP position | Genotypes of individual 4 | | Genotypes of individual 5 | | Genotypes of individual 6 | |
|---|---|---|---|---|---|---|
| 734 | **T** | A | T | **T** | A | A |
| 957 | **C** | G | G | **C** | C | G |

**Table S8.** A population-specific reference panel of individual genotypes, sharing the same ancestral population as homolog $g$. The reference panel describes the genotypes of 3 individual at SNP positions, with the haplotype that was observed in reads **C** and **E** being highlighted.

where $n$ is the number of sampled reads, $(\mathbb{Z}_2)^2 = \mathbb{Z}_2 \times \mathbb{Z}_2 = \big\{(a,b) \mid a \in \{0,1\} \ \text{ and } \ b \in \{0,1\}\big\}$ is a 2-fold Cartesian product and similarly $(\mathbb{Z}_2)^n = \prod_{i=1}^{n} \mathbb{Z}_2$ is a $n$-fold Cartesian product of $\mathbb{Z}_2$. Thus, we sum over all the possible configurations of reads, using the "occupation-number" representation. In addition, XOR is the logical operation "Exclusive or", e.g, (0,1,0) XOR (1,1,1)=(1,0,1). The rationale behind the XOR operation is that every read that is not associated with homolog $f$ is necessarily associated with homolog $g$.

We notice that when both homolog $f$ and homolog $g$ are associated with the same ancestral population, every term in the sum appears twice. Hence, we simplify the expression by summing over half of the terms and multiply the result by 2:

$$P(\mathbf{A} \wedge \mathbf{B} \ldots) = \frac{1}{2^n} \sum_{x \in (\mathbb{Z}_2)^n} f\big(x\big) f\big(x \text{ XOR } (1,1,\ldots,1)\big) = \frac{1}{2^{n-1}} \sum_{x \in \mathbb{Z}_1 \times (\mathbb{Z}_2)^{n-1}} f\big(x\big) f\big(x \text{ XOR } (1,1,\ldots,1)\big), \tag{21}$$

where $\mathbb{Z}_1 \times (\mathbb{Z}_2)^2 = \big\{(0,a,b) \mid a \in \{0,1\} \ \text{ and } \ b \in \{0,1\}\big\}$, meaning that we sum over all the sequences of length $n$ with the first element fixed to 0.

#### The "matched" and "unmatched" statistical models for "no admixture" and "distant admixture" scenarios.

The statistical models of the matched haplotypes hypothesis as well as the unmatched haplotypes hypothesis for individuals with non-admixed ancestry with between two to four sampled reads are given below:

##### 1+1 Reads

$$P^{\text{non-admixed}}_{\text{unmatched}}(A \wedge B) = f(A)f(B)$$

$$P^{\text{non-admixed}}_{\text{matched}}(A \wedge B) = \frac{1}{2}\left[g(B) + g(\varnothing)f(B)\right]$$

##### 1+2 Reads

$$P^{\text{non-admixed}}_{\text{unmatched}}(A \wedge B \wedge C) = \frac{1}{2}f(A)\left[f(BC) + f(B)f(C)\right]$$

$$P^{\text{non-admixed}}_{\text{matched}}(A \wedge B \wedge C) = \frac{1}{4}\left[f(B)g(C) + f(C)g(B) + f(BC)g(\varnothing) + g(BC)\right]$$

##### 1+3 Reads

$$P^{\text{non-admixed}}_{\text{unmatched}}(A \wedge B \wedge C \wedge D) = \frac{1}{4}f(A)\left[f(BCD) + f(BC)f(D) + f(BD)f(C) + f(CD)f(B)\right]$$

$$P^{\text{non-admixed}}_{\text{matched}}(A \wedge B \wedge C \wedge D) = \frac{1}{8}\left[f(BCD)g(\varnothing) + f(B)g(CD) + f(C)g(BD) + f(D)g(BC)\right] + (f \leftrightarrow g)$$

##### 1+4 Reads

$$P^{\text{non-admixed}}_{\text{unmatched}}(A \wedge B \wedge C \wedge D \wedge E) = \frac{1}{8}f(A)\left[f(B)f(CDE) + f(C)f(BDE) + f(D)f(BCE) + f(E)f(BCD) + f(BE)f(CD) + f(CE)f(BD) + f(DE)f(BC) + f(BCDE)\right]$$

$$P^{\text{non-admixed}}_{\text{matched}}(A \wedge B \wedge C \wedge D \wedge E) = \frac{1}{16}\left[f(B)g(CDE) + f(C)g(BDE) + f(D)g(BCE) + f(E)g(BCD) + f(BE)g(CD) + f(CE)g(BD) + f(DE)g(BC) + f(BCDE)g(\varnothing)\right] + (f \leftrightarrow g)$$

- The function $f$ is a joint frequency distribution that is derived from a reference panel. The reads $A, B, C$ and $D$ are represented by vectors of chromosomal positions and nucleotides. For brevity we use the notation $f(ABC \ldots Z) \equiv f(A, B, C, \ldots, Z)$, i.e. when $AB$ appears as an argument of a function it reflects two distinct arguments, $A$ and $B$ and not a scalar product of the two.
- In all the models read $A$ is drawn from the DNA library of the monosomy, while the rest of the reads are drawn from the DNA library of the disomy.
- The "unmatched" model of $1 + n$-reads is merely the disomy model of $n$-reads for non-admixed multiplied by the joint frequency $f(A)$; More information about the disomy model for non-admixed can found in Ariad et al. [28].
- The "matched" model of $1 + n$-reads is the disomy model of $n$-reads for recent-admixture with effective joint frequency distributions. The first joint frequency distribution is associated with the "matched" haplotypes, while the second is associated with the "unmatched" haplotypes. Therefore, we define the distribution of the "matched" haplotypes as

  $$g(BC \ldots Z) \equiv f(ABC \ldots Z),$$

  where read $A$ is drawn from the library of the monosomy, while the rest of the reads are drawn from the library of the disomy. The joint frequency distribution of the "unmatched" haplotypes, denoted as $f$, was derived from a reference panel of a population. In addition, we define $g(\varnothing) \equiv f(A)$ and $f(\varnothing) \equiv 1$ because read $A$ is present in all the terms of a "matched" model. For more information about the disomy model for recent-admixture see [28])

- The notation $(f \leftrightarrow g)$ is used to represent the sum of all the other terms in the expression with $f$ and $g$ exchanged, e.g., $f(A)g(B) + (f \leftrightarrow g) = f(A)g(B) + g(A)f(B)$.
- For distant-admixtures we define an effective frequency,

$$f(x) \equiv \alpha_1 f_1(x) + \alpha_2 f_2(x) + \ldots + \alpha_n f_n(x),$$

which takes into account that a haplotype have originated from one of several populations with probability $\alpha_i$ to originate from the $i$th population.

- Since all the reads that were drawn from the library of the monosomy originated from the same DNA molecule, we can easily generalize the models to any number of reads from the monosomy library by the substitution $A \to A_1, A_2, \ldots$. For example, the models for $2 + 1$ reads is

$$P^{\text{non-admixed}}_{\text{unmatched}}(A_1 \wedge A_2 \wedge B) = f(A_1 A_2) f(B)$$

$$P^{\text{non-admixed}}_{\text{matched}}(A_1 \wedge A_2 \wedge B) = \frac{1}{2}\big[g(B) + g(\varnothing) f(B)\big],$$

where we redefine $g(BC \ldots Z) \equiv f(A_1 A_2 BC \ldots Z)$.

#### The "matched" and "unmatched" statistical models for admixture in the previous generation

Here we consider an individual descended from parents of distinct ancestries (hereafter termed "recent admixed"). The "matched" and "unmatched" statistical models for such recent admixed individuals for between two to four reads are given below:

##### 1+1 Reads

$$P^{\text{recent-admixed}}_{\text{unmatched}}(A \wedge B) = \frac{1}{4}\left[f_1(A) + f_2(A)\right]\left[f_1(B) + f_2(B)\right]$$

$$P^{\text{recent-admixed}}_{\text{matched}}(A \wedge B) = \frac{1}{4}\left[g_1(B) + f_2(B)g_1(\varnothing) + f_1(B)g_2(\varnothing) + g_2(B)\right]$$

##### 1+2 Reads

$$P^{\text{recent-admixed}}_{\text{unmatched}}(A \wedge B) = \frac{1}{8}\left[f_1(A) + f_2(A)\right]\left[f_1(BC) + f_2(BC) + f_1(B)f_2(C) + f_1(C)f_2(B)\right]$$

$$\begin{aligned} P^{\text{recent-admixed}}_{\text{matched}}(A \wedge B) =& \frac{1}{8}\left[f_1(B)g_2(C) + f_1(C)g_2(B) + f_1(BC)g_2(\varnothing) + f_1(\varnothing)g_2(BC)\right] + \\ & \frac{1}{8}\left[g_1(B)f_2(C) + g_1(C)f_2(B) + g_1(BC) + g_1(\varnothing)f_2(BC)\right] \end{aligned}$$

##### 1+3 reads

$$P^{\text{recent-admixed}}_{\text{unmatched}}(A \wedge B \wedge C \wedge D) = \frac{1}{16}\left[f_1(A) + f_2(A)\right]\left[f_1(B)f_2(CD) + f_1(C)f_2(BD) + f_1(D)f_2(BC) + f_1(BCD)\right] + (f_1 \leftrightarrow f_2)$$

$$\begin{aligned} P^{\text{recent-admixed}}_{\text{matched}}(A \wedge B \wedge C \wedge D) =& \frac{1}{16}\left[f_1(B)g_2(CD) + f_1(C)g_2(BD) + f_1(D)g_2(BC) + f_1(BCD)g_2(\varnothing) + (f_1 \leftrightarrow g_2)\right] + \\ & \frac{1}{16}\left[g_1(B)f_2(CD) + g_1(C)f_2(BD) + g_1(D)f_2(BC) + g_1(BCD)f_2(\varnothing) + (f_2 \leftrightarrow g_1)\right] \end{aligned}$$

##### 1+4 reads

$$\begin{aligned} P^{\text{recent-admixed}}_{\text{unmatched}}(A \wedge B \wedge C \wedge D \wedge E) =& \frac{1}{32}\left[f_1(A) + f_2(A)\right]\left[f_1(B)f_2(CDE) + f_1(C)f_2(BDE) + f_1(D)f_2(BCE) + f_1(E)f_2(BCD) + \right. \\ & \left. f_1(BE)f_2(CD) + f_1(CE)f_2(BD) + f_1(DE)f_2(BC) + f_1(BCDE) + (f_1 \leftrightarrow f_2)\right] \end{aligned}$$

$$\begin{aligned} P^{\text{recent-admixed}}_{\text{matched}}(A \wedge B \wedge C \wedge D \wedge E) =& \frac{1}{32}\left[f_1(E)g_2(BCD) + f_1(B)g_2(CDE) + f_1(C)g_2(BDE) + f_1(D)g_2(BCE) + \right. \\ & \left. f_1(BE)g_2(CD) + f_1(CE)g_2(BD) + f_1(DE)g_2(BC) + f_1(BCDE)g_2(\varnothing) + (f_1 \leftrightarrow g_2)\right] + \\ & \frac{1}{32}\left[g_1(E)f_2(BCD) + g_1(B)f_2(CDE) + g_1(C)f_2(BDE) + g_1(D)f_2(BCE) + \right. \\ & \left. g_1(BE)f_2(CD) + g_1(CE)f_2(BD) + g_1(DE)f_2(BC) + g_1(BCDE)f_2(\varnothing) + (g_1 \leftrightarrow f_2)\right] \end{aligned}$$

- The function $f_i$ is a joint frequency distribution that is derived from a reference panel of the $i$ population. The reads $A, B, C$ and $D$ are represented by vectors of chromosomal positions and nucleotides. For brevity we use the notation $f_i(ABC \dots Z) \equiv f_i(A, B, C, \dots, Z)$, i.e. when $AB$ appears as an argument of a function it reflects two distinct arguments, $A$ and $B$ and not a scalar product of the two.
- In all the models read $A$ is drawn from the DNA library of the monosomy, while the rest of the reads are drawn from the DNA library of the disomy.
- The "unmatched" model of $1 + n$-reads is merely the disomy model of $n$-reads for recent-admixtures multiplied by the term $[f_1(A) + f_2(A)]/2$. This term takes into account that the unmatched haplotypes can be associated with one of two populations with equal probability. More information about the disomy model for recent-admixtures can found in [28].

- The "matched" model of $1+n$-reads is based on a disomy model of $n$-reads for recent-admixture. The first joint frequency distribution is associated with the "matched" haplotypes, while the second is associated with the "unmatched" haplotypes. Therefore, we define the distribution of the "matched" haplotypes as

$$g_i(BC\ldots Z) \equiv f_i(ABC\ldots Z),$$

  where read $A$ is drawn from the library of the monosomy, while the rest of the reads are drawn from the library of the disomy. The joint frequency distribution of the "unmatched" haplotypes, denoted as $f_i$, was derived from a reference panel of population $i$ and $i = 1, 2$. In addition, we define $g_i(\varnothing) \equiv f_i(A)$ and $f_i(\varnothing) \equiv 1$ because read $A$ should be present in all the terms of the statistical models. For more information about the disomy model for recent-admixture see [28])

- The notation $(f_1 \leftrightarrow f_2)$ is used to represent the sum of all the other terms in the expression with $f_1$ and $f_2$ exchanged, e.g., $f_1(A)f_2(B) + (1 \leftrightarrow 2) = f_1(A)f_2(B) + f_2(A)f_1(B)$.

- Since all the reads that were drawn from the library of the monosomy originated from the same DNA molecule, we can easily generalize the models to any number of reads from the monosomy library by the substitution $A \to A_1, A_2, \ldots$. For example, the models for $2+1$ reads is

$$P^{\text{recent-admixed}}_{\text{unmatched}}(A_1 \wedge A_2 \wedge B) = \frac{1}{4}\left[f_1(A_1A_2) + f_2(A_1A_2)\right]\left[f_1(B) + f_2(B)\right]$$

$$P^{\text{recent-admixed}}_{\text{matched}}(A \wedge B) = \frac{1}{4}\left[g_1(B) + f_2(B)g_1(\varnothing) + f_1(B)g_2(\varnothing) + g_2(B)\right]$$

  where we redefine $g_i(BC\ldots Z) \equiv f_i(A_1A_2BC\ldots Z)$.

#### Simulating a parental gamete and a diploid offspring in distant admixture scenarios.

First, we divide the genome into regions of 1 - 10 Mbp. For simplicity we assume a distant admixture that involves only two ancestral populations, but it is straight forward to extend the generative model to any number of ancestral populations. We choose an ancestral makeup where $\%(100 \cdot \alpha)$ of the haplotypes are associated with population A, while $\%\big(100 \cdot (1-\alpha)\big)$ of the haplotypes are associated with population B. Then, we draw 3 pairs of effective haploids with each pair composed of one haploid that is associated with population A and a second one that is associated with population B.

We consider the first pair and to each genomic region we assign either the effective haploid from population A with a probability $\alpha$ or the effective haploid from population B with a probability $1-\alpha$. Then, we repeat the process for the two other pairs, so that three effective haploids would be assigned to each genomic region. In each region, the first two haploids would be used to simulate the diploid offspring, while the first and third haploids would be used to simulate the parental gamete and the unrelated gamete, respectively. Then, we simulate reads by selecting a random position along the chromosome from a uniform distribution, representing the midpoint of an aligned read with a given length. Based on the selected position, one out of the six haplotypes was drawn from a discrete distribution,

$$f(h,x) = \begin{cases} p_{1,A}(x) & , h = 1A \\ p_{1,B}(x) & , h = 1B \\ p_{2,A}(x) & , h = 2A \\ p_{2,B}(x) & , h = 2B \\ p_{3,A}(x) & , h = 3A \\ p_{3,B}(x) & , h = 3B \end{cases}, \tag{22}$$

where the probability of haplotype $h$ depends on the position of the read, $x$.

In each genomic region either $p_{i,A} = 0$ or $p_{i,B} = 0$ with $i = 1, 2, 3$, and hence within each genomic region the discrete distribution reduces to

$$f(h) = \begin{cases} p_1 & , h = 1 \\ p_2 & , h = 2 \\ p_3 & , h = 3 \end{cases}. \tag{23}$$

When simulating a diploid offspring, the first haplotype is just as likely as the second haplotype, $p_2 = p_1$ and the third haplotype is absent, $p_3 = 0$. Similarly, for the parental gamete $p_1 = 1$ and $p_2 = p_3 = 0$, while for the unrelated gamete $p_3 = 1$ and $p_1 = p_2 = 0$. Then, from the selected haplotype, $h$, a segment of length $l$ that is centered at the selected chromosomal position, $x$, is added to simulated data, mimicking the process of short-read sequencing. This process of generating simulated sequencing data is repeated until the desired depth of coverage is attained.

| | Ancestral population A/B | | |
|---|---|---|---|
| Regions | Pair 1 | Pair 2 | Pair 3 |
| 1 | 1A | 2B | 3A |
| 2 | 1B | 2A | 3A |
| 3 | 1B | 2B | 3B |
| 4 | 1A | 2B | 3A |
| 5 | 1B | 2A | 3B |
| 6 | 1A | 2A | 3B |

**Table S9.** We assign 3 haploids to each region in the genome. Each haploid is selected from a pair of haploids, where each haploid in the pair is associated with a different ancestral population. The probability to draw a haploid that is associated with population A is $\alpha$, while it is $1-\alpha$ for population B. In the table we show a possible outcome of this procedure for 6 regions. Haploids that were drawn from the first two pairs are used to simulate a diploid offspring, while haploids that were drawn from the first and third pair are used to simulate the matched and unmatched gametes, respectively.

#### An example for calculating the score of a read

Here we use the score metric that was introduced in the subsection "Prioritizing informative reads" to calculate the score of an example read, shown Figure S23. In this example, we consider the ancestral makeup: 30% population A and 70% population B. In addition, we assume that for each population we have a suitable reference panel.

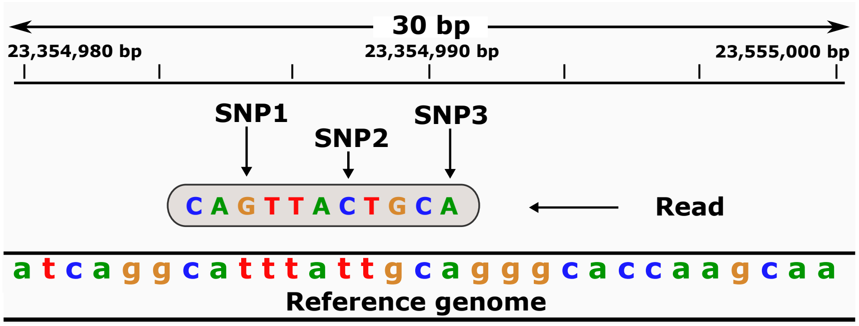


| Haplotype | Joint Frequency A | Joint Frequency B | Effective Frequency | Score Increment $= \begin{cases} 1, & 0.1 < f_{\text{eff}} < 0.9 \\ 0, & \text{otherwise} \end{cases}$ |
|---|---|---|---|---|
| $GTC$ | 0.22 | 0.13 | $0.3 * 0.22 + 0.7 * 0.13 = 0.157$ | +1 |
| $GTA$ | 0.02 | 0.07 | 0.055 | 0 |
| $GCA$ | 0.15 | 0.17 | 0.164 | +1 |
| $GCC$ | 0.28 | 0.36 | 0.336 | +1 |
| $TTC$ | 0.01 | 0.12 | 0.087 | 0 |
| $TTA$ | 0.37 | 0.21 | 0.258 | +1 |
| $TCA$ | 0.03 | 0.11 | 0.086 | 0 |
| $TCC$ | 0.01 | 0.07 | 0.052 | 0 |
| | | | | Total score: 4 |

**Figure S23.** (Top) A read that is aligned to a reference genome. (Bottom) A table of possible haplotypes in the chromosomal region that overlaps with the read, the frequencies of the haplotypes (as estimated from two reference panels), and the scores of the haplotypes.

#### Estimating the probability of correctly calling a crossover

Let us consider a crossover that was identified via the procedure in the section " Identifying chromosomal crossovers". A crossover is characterized as a transition between a region of matched haplotypes and a region of unmatched haplotypes upon comparison of sibling embryos. In each region, we cluster all the genomic windows into a bin by aggregating the mean and the variance of LLRs in each genomic window, as described in the section "Aggregating log likelihood ratios across consecutive windows". For a given chromosome, we denote the nearest bin to the left (right) of the $n$th crossover as bin $n, L$ (bin $n, R$).

Following the principles of Mendelian inheritance, we assume that probability of having a region with unmatched or matched haplotypes is half, $P(U) = P(M) = \frac{1}{2}$ with $U$ and $M$ denoting the unmatched and matched haplotypes, respectively.

In addition, we define $P(U|M)$ as the probability of calling a bin "unmatched" given that it is actually "matched". The relations between the four possible conditional probabilities and the diagnostic ability of the classifier are:

$$P(U|U) = \text{TPR}, \quad P(U|M) = \text{FNR}, \quad P(M|U) = \text{FPR}, \quad P(M|M) = \text{TNR},$$

where TPR is the true positive rate, FNR is the false negative rate, FPR is the false negative rate and TNR is the true negative rate.

Now let us consider two adjacent bins, where bin $n, L$ is predicted as "unmatched" and bin $n, R$ as "matched". This switch from "unmatched" to "matched" indicates the presence of a crossover with a probability of

$$P(U, M) = P(U, M|U, M)P(U, M) = \frac{1}{4}P(U|U)P(M|M) = \frac{1}{4}\text{TPR}_{n,L} \cdot \text{TNR}_{n,R},$$

where $\text{TPR}_{n,L}$ is the true positive rate of the classifier for bin $n, L$. Similarly the probability of correctly calling a switch from "matched" to "unmatched" is

$$P(M, U) = P(M, U|M, U)P(M, U) = \frac{1}{4}P(M|M)P(U|U) = \frac{1}{4}\text{TNR}_{n,L} \cdot \text{TPR}_{n,R}.$$

Next, we consider the probability of missing a crossover because two adjacent bins are predicted to be "unmatched":

$$P(U, U|U, M)P(U, M) + P(U, U|M, U)P(M, U) = \frac{1}{4}\left(P(U|U)P(U|M) + P(U|M)P(U|U)\right) = \frac{1}{4}\left(\text{TNR}_{n,L} \cdot \text{FNR}_{n,R} + \text{FNR}_{n,L} \cdot \text{TNR}_{n,R}\right),$$

Similarly, the probability of missing a crossover because two adjacent bins are predicted to be "matched" by the classifier is

$$P(M, M|M, U)P(M, U) + P(M, M|U, M)P(U, M) = \frac{1}{4}\left(P(M|M)P(M|U) + P(M|U)P(M|M)\right) = \frac{1}{4}\left(\text{TPR}_{n,L} \cdot \text{FPR}_{n,R} + \text{FPR}_{n,L} \cdot \text{TPR}_{n,R}\right).$$
